# A Fibrous Optofluidic Probe for Endovascular Sensing of Blood Oxygenation and Flow Velocity

**DOI:** 10.64898/2026.11.03.763613

**Authors:** Bozhen Zhang, Xianli Lv, Zhongyin Ye, Xin Wang, Weiming Kong, Huachen Zhang, Derun Li, Jiazheng Chen, Zican Xie, Dawid Sheng, Xue Cai, Yuqi Wang, Qianyong Yang, He Ding, Lan Yin, Xing Sheng

**Affiliations:** School of Materials Science and Engineering, The Key Laboratory of Advanced Materials of Ministry of Education, State Key Laboratory of Flexible Electronics Technology, Tsinghua University, Beijing, 100084, China; Department of Neurosurgery, Beijing Tsinghua Changgung Hospital, School of Clinical Medicine, Tsinghua Medicine, Tsinghua University, Beijing, 102218, China; Department of Electronic Engineering, Beijing National Research Center for Information Science and Technology, Institute for Precision Medicine, State Key Laboratory of Flexible Electronics Technology, Beijing Key Laboratory of Clinical Innovation and Translation for Active Implantable and Interventional Medical Devices, IDG/McGovern Institute for Brain Research, Tsinghua University, Tsinghua University, Beijing, 100084, China; Institute of Optical Information Science and Technology, School of Physical Science and Engineering, Beijing Jiaotong University, Beijing, 100044, China; Jiangxi Provincial Key Laboratory of Cell Precision Therapy, School of Basic Medical Sciences, Jiujiang University, Jiujiang, 332005, China; Beijing Engineering Research Center of Mixed Reality and Advanced Display, School of Optics and Photonics, Beijing Institute of Technology, Beijing, 100081, China

## Abstract

Local hemodynamic information including blood oxygenation and flow velocity could guide vascular interventions, yet obtaining these parameters in deep, narrow vessels remains challenging. Here, we report a flexible, submillimeter-scale fibrous optofluidic probe for endovascular sensing of blood oxygenation and flow velocity. The probe integrates a polymer optical fiber for light transmission with a microfluidic tube for dye injection, enabling spectroscopic oximetry and fluorescence dilution-based velocimetry through a single device. Via vascular intervention, we demonstrate local hemodynamic sensing in rabbit arteries and angiography-guided deployment in canine carotid arteries. The probe captures local changes in blood oxygenation and flow velocity during vascular occlusion and reperfusion, revealing hemodynamic disturbances that are not reflected in peripheral monitors. These results illustrate the complementary information provided by local oximetry and velocimetry and establish a compact optofluidic platform for quantitative hemodynamic monitoring during vascular interventions.

## INTRODUCTION

Cardiovascular and cerebrovascular diseases (e.g., heart attacks and stroke) are major causes of death and disability worldwide^1,2^. Endovascular intervention is a rapidly expanding minimally invasive therapeutic strategy for these conditions^3–5^. During these procedures, digital subtraction angiography (DSA) guides the navigation and deployment of therapeutic devices such as stents^6^, coils^7^, and flow diverters^8^ through the vasculature to the region of interest to treat disorders including aneurysms^9^ and arterial stenoses^10^. However, procedural disturbances such as vasospasm, thrombosis, and unintended vascular occlusion can compromise local blood flow and oxygen delivery^11–13^, complicating intraoperative assessment of the procedural result. Although angiography provides essential anatomical and flow-related information, continuous, quantitative assessment of local hemodynamics remains difficult^14–15^. Measurements obtained directly at the intervention site could complement imaging by revealing functional changes during treatment and recovery.

To enable earlier and more reliable detection, intraoperative quantitative monitoring techniques of local blood oxygenation and flow velocity are being actively pursued. Measuring these parameters could help determine the origin of hemodynamic changes, distinguishing shifts in blood oxygenation from disturbances in flow during vascular occlusion and reperfusion. Existing monitoring approaches address different aspects of this need, including Doppler flowmetry^16^, pulse oximetry^17,18^, intravascular ultrasound^19,20^, oximetric catheters^21,22^, yet each provides valuable but inherently limited information. For instance, non-invasive ultrasonic and optical devices suffer from significant tissue scattering and absorption, offering limited accuracy particularly when interrogating intracranial and other deep-tissue vessels^23,24^. Implantable extravascular devices circumvent these constraints but require surgical exposure for placement, restricting their use to accessible or superficial vascular sites. Intravascular sensors are well suited to interventional delivery, but most remain centimeter-scale, limiting their use to larger vessels, and typically measure only a single parameter^25–27^. Integrating oxygenation and velocity measurements within a single miniature device would therefore provide a useful complement to existing interventional monitoring.

Here, we develop a flexible, submillimeter fibrous optofluidic probe (FOP) (Fig. 1a) for endovascular sensing of blood oxygenation and flow velocity. The FOP sensor integrates a polymeric optical fiber for light transmission within a microfluidic tube for dye injection, enabling spectroscopic oximetry and fluorescence dilution-based velocimetry. This compact optofluidic design allows for reliable operation within deep, narrow vasculature. We characterize its sensing performance, demonstrate local hemodynamic measurements in rabbit arteries, and establish angiography-guided deployment in canine carotid arteries. The FOP continuously captures local hemodynamic changes, detecting flow compromise and impaired oxygen delivery associated with varied hemodynamic conditions. These results demonstrate the complementary information provided by dual-parameter optofluidic sensing and support its potential for quantitative hemodynamic monitoring during vascular interventions.

**Figure 1.**
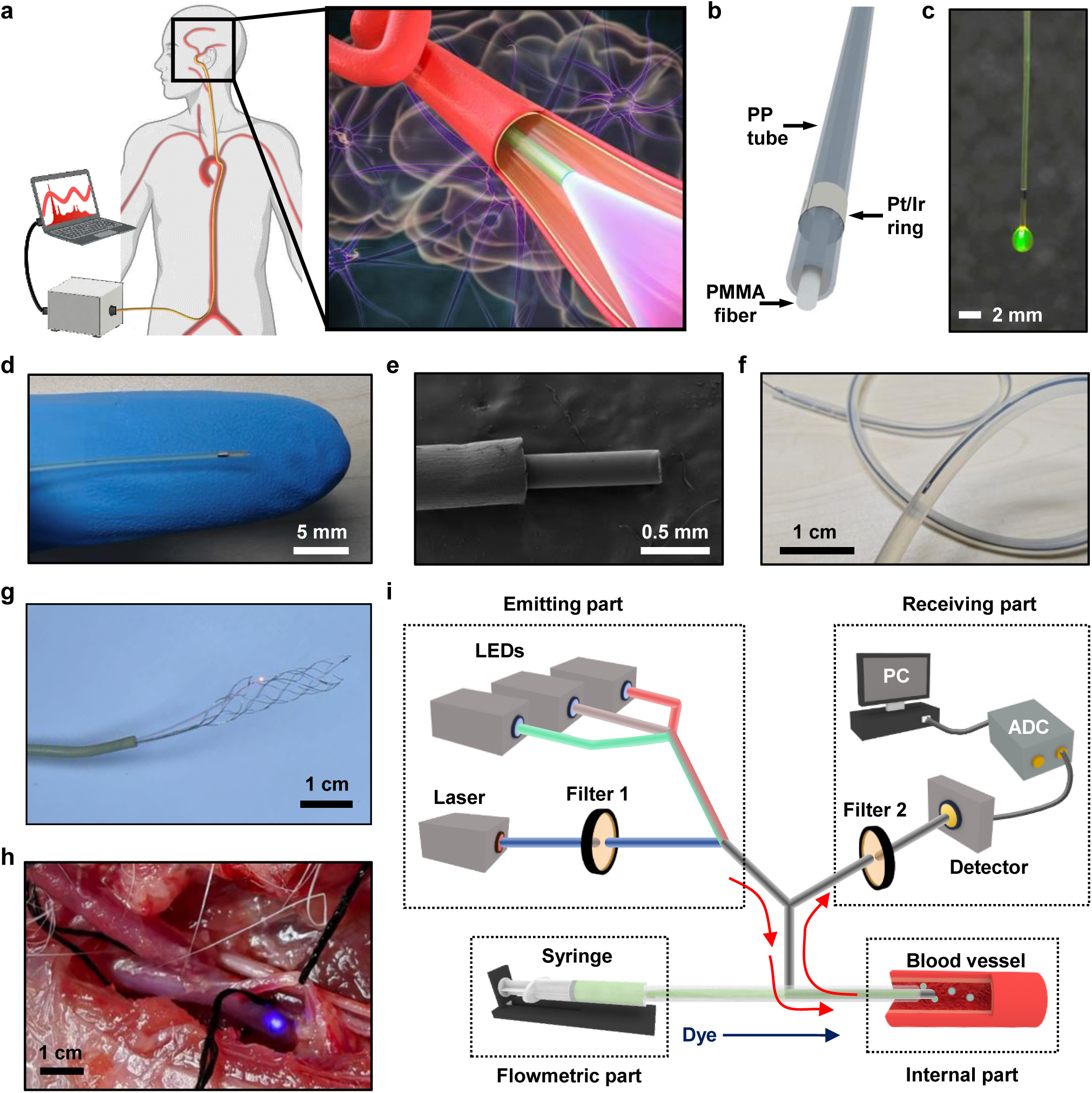
Concept and design of the fibrous optofluidic probe (FOP) sensor for blood oxygenation and blood velocity monitoring in the vessel. **a.** Schematic illustration of the FOP sensor designed for intraoperative monitoring of vascular intervention surgeries in thin vasculature in deep tissue (e.g., cerebral arteries). **b.** Structure of the FOP sensor, including a PMMA plastic optical fiber for light transmission, a PP microfluidic channel for solution injection, and a Pt/Ir ring as a radiopaque marker. **c.** Image of the sensor tip with a droplet of injected dye lightened by the optical fiber. **d.** Photo and **e.** electron microscopic image of the probe tip, with an outer diameter of around 0.5 mm. **f.** Image of a flexible FOP sensor inserted in a thin curvy tube. The microfluidic channel in the FOP is filled with ink for visualization. **g.** Image of an FOP sensor and a stent-holding guidewire through a 6-Fr catheter (outer diameter of 1.67 mm), showing its compatibility with interventional systems for clinical use. **h.** Image of a sensor implanted in a rabbit femoral artery (inner diameter of ∼2 mm). **i.** Scheme of the customized sensing system, including light emitting and receiving parts, solution injection, endovascular transduction, and data acquisition components.

## RESULTS

### Materials and design of the FOP sensor

The FOP sensing system comprises two main components: the biocompatible, flexible endovascular probe and the extracorporeal optofluidic system (Fig. 1a). The miniaturized probe (Fig. 1b and Fig. S1) consists of a polymethyl methacrylate (PMMA) multimode optical fiber (diameter of ∼250 μm) and a polypropylene (PP) microfluidic tube (inner diameter of ∼400 μm; outer diameter of ∼500 μm). The PMMA fiber serves as the waveguide for optical transmission across the visible and near-infrared spectrum (Fig. S2), while the PP tube acts as a protective sheath that prevents scratching during storage and implantation. Additionally, the annular space between the tube and the fiber forms a microfluidic channel for delivering dye solution into the bloodstream (Fig. 1c and Movie S1). A platinum/iridium (Pt/Ir) ring (outer diameter of ∼610 μm) is affixed to the probe tip as a radiopaque marker, rendering the device visible under angiography. The overall probe diameter is submillimeter (∼500 μm), comparable to clinically used guidewires (0.35–0.6 mm^28–30^) and substantially smaller than major cerebral, coronary, and peripheral arteries (1.5–4 mm^31–33^) (Figs. 1d and 1e).

Mechanical characterization reveals that the FOP exhibits an effective modulus of 2.12 GPa, with a bending stiffness of 4.23 N·mm^2^ and an elastic limit of 1.29% (Fig. S3). The FOP exhibits transmission losses of 0.41–1.84 dB/m across the 470–810 nm wavelength range and remains insensitive to mild bending in curved vessels (Fig. S4). These mechanical and optical properties are comparable to, or even surpass, those of clinically used guidewires^34,35^ and previously reported implantable optical fibers^36–40^ (Table S1), support its optical sensing and flexible deployment within narrow vascular geometries. Owing to its flexibility, the FOP can be navigated into tortuous, branching lumens without guidewire assistance (Fig. 1f and Movie S2). Moreover, the probe can be deployed alongside a stent-holding guidewire through a clinical microcatheter (Fig. 1g). As a functional demonstration, Fig. 1h and Movie S3 show the light-emitting FOP inserted into a rabbit femoral artery, confirming its operability within narrow-caliber vasculature.

We further develop an extracorporeal optofluidic system to support endovascular hemodynamic monitoring of the FOP sensor (Fig. 1i, Fig. S5 and Table S2). The optical subsystem uses a 2×1 fiber-optic coupler to establish a two-way light path: light from multiple LEDs and laser sources is delivered into the FOP, while reflected or emitted light from the blood is collected by a photodetector and recorded via an analog-to-digital converter. The fluidic subsystem comprises a programmable syringe pump that delivers dye boluses for velocimetry through the microfluidic channel. A customized three-way connector (Figs. S1c and S1d) interfaces the extracorporeal system with the optical fiber and microfluidic tube of the FOP, enabling independent and interference-free optical and fluidic operation. This configuration supports a range of optical sensing modalities, including intensity-, spectrum-, and lifetime-based measurements. In the present work, we integrate spectroscopic oximetry and dilution-based velocimetry within this framework to achieve real-time hemodynamic monitoring during interventional procedures.

### FOP-based oximetric sensing

We implement spectroscopic oximetry based on the distinct absorption spectra of oxygenated hemoglobin (HbO_2_) and deoxygenated hemoglobin (Hb) (Figs. 2a and 2b). Blood oxygen saturation (SO_2_), which is the fraction of HbO_2_ in the total hemoglobin (HbO_2_ and Hb) concentration, is derived from the relative concentrations of these two chromophores. Three wavelengths, 530 nm (green), 660 nm (red), and 810 nm (infrared), are selected for their complementary sensitivities. The 660 nm wavelength lies at the maximum differential absorption between HbO_2_ and Hb, providing high sensitivity to SO_2_. The 530 nm and 810 nm wavelengths are near the isosbestic points, rendering them primarily sensitive to total hemoglobin concentration rather than oxygenation state^41–43^. The three LEDs are sequentially pulsed under microcontroller control, and the temporally resolved photodetector signal is used to extract spectral information for SO_2_ calculation (Fig. S6, Table S3 and Analysis S1).

**Figure 2.**
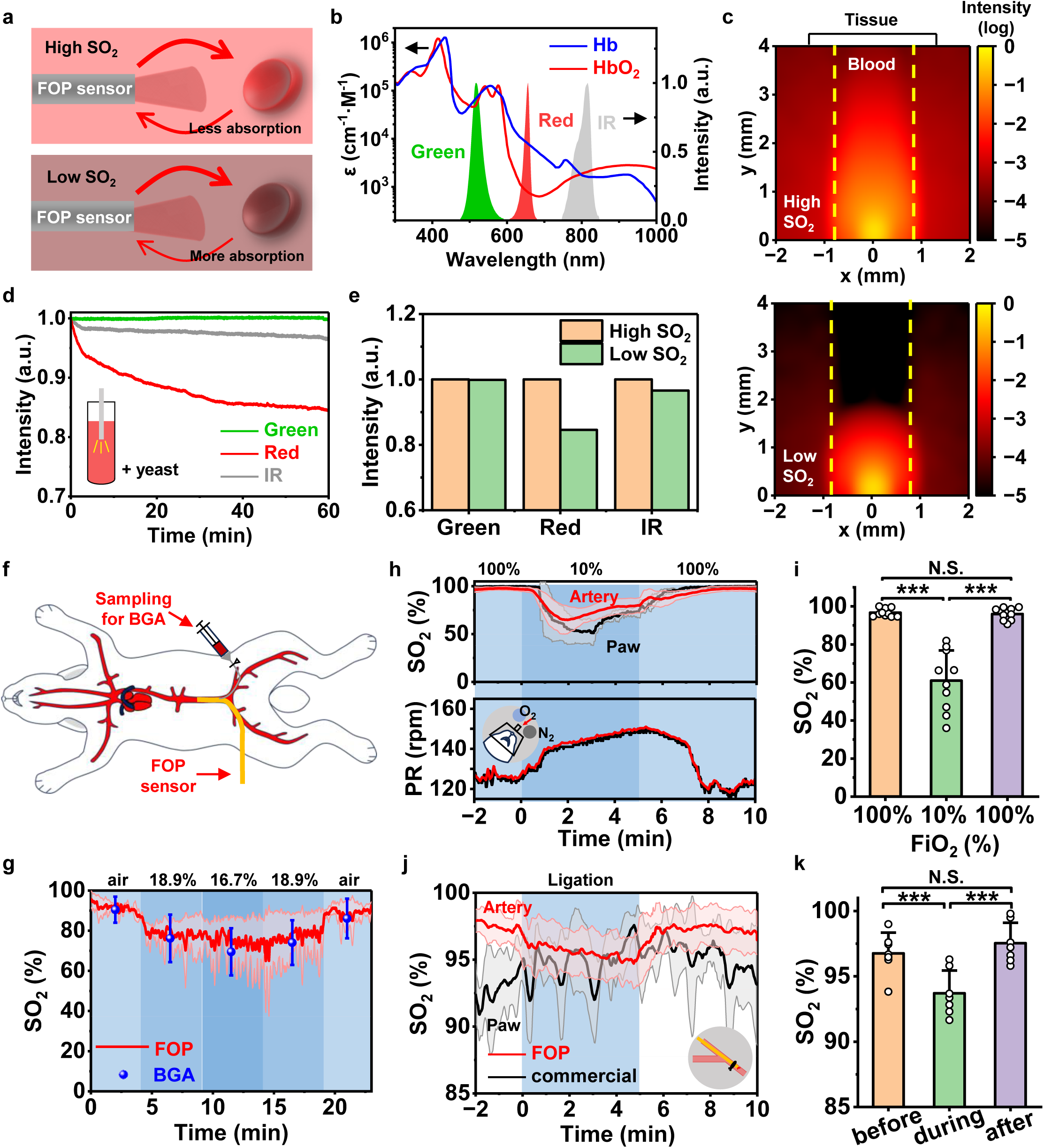
FOP-based blood oximetry. **a.** Principle of blood oxygen saturation (SO_2_) sensing. Variation of SO_2_ causes different light absorption, resulting in different intensities of receiving light recorded by the fiber. **b.** Absorption spectra of Hb and HbO_2_, as well as emitting spectra of green, red and IR LEDs used for SO_2_ sensing. **c.** Numerical simulations showing power distribution of 660 nm light emitting into a blood vessel with high SO_2_ (98%, upper) and low SO_2_ (0%, lower) blood. **d.** Measured light intensity change in the porcine blood, in which SO_2_ is regulated by adding yeast. **e.** Comparison of light intensity measured at different wavelengths before and after oxygen consumption by yeast. **f.** Illustration of in vivo SO_2_ sensing experiment operated in a rabbit abdominal aorta model. **g.** Dynamic SO_2_ sensing results measured at different fractions of inspired O_2_ (*F*_iO2_) by our FOP sensor, compared with data simultaneously collected by a commercial blood gas analyzer (BGA) (3 trials, *n* = 2 rabbits). **h.** Simultaneously measured SO_2_ and pulse rate (PR) change at different FiO_2_. Red: results detected by our FOP sensor within the femoral artery; Black: results collected externally by a commercial pulse oximeter mounted at the paw. **i.** Comparison of SO_2_ results at different FiO_2_ (10 trials, *n* = 3 rabbits). **j.** Measured SO_2_ change when acute ischemia and reperfusion are created by ligation in the upstream artery. **k.** Comparison of SO_2_ results in the artery ligation model (7 trials, *n* = 4 rabbits). Paired two-tailed *t*-tests. \**p* < 0.05, \*\**p* < 0.01, \*\*\**p* < 0.001, N.S. *p* > 0.05. Data are presented as mean ± S.D.

We first evaluate the feasibility of this fiber-based oximetry through numerical simulation. Modelled results in an artificial vessel structure (Fig. 2c and Fig. S7) show that red-light absorption is highly sensitive to the change of SO_2_, with the recorded intensity decreasing by ∼49% when SO_2_ drops from the high level (98%) to the low level (0%), whereas green and infrared intensities decrease by only 5.2% and 17.0%, respectively. We then immerse the FOP sensor in the whole blood and monitored signal changes as yeast respiration progressively reduced SO_2_ (Fig. 2d). The red-light intensity decreases by 15.4%, whereas green and infrared signals show much smaller drops (0.1% and 3.4%, respectively) (Fig. 2e). While these trends are qualitatively consistent with the simulation, the absolute intensity changes are smaller. This discrepancy is attributed to two factors: background reflection in the fiber-optic system introduces a static light component unrelated to blood oxygenation, and the initial SO_2_ in blood equilibrated with air is below the simulated high level (98%), reducing the magnitude of the subsequent decline. Collectively, these data confirm that the FOP sensor responds sensitively to SO_2_ changes in blood.

To establish the model relating recorded light intensity to SO_2_, we perform in vivo calibration using a blood gas analyzer (BGA) as the reference standard^44^. In a rabbit model, the FOP sensor is implanted in the abdominal aorta, where it continuously record optical signals while BGA measurements are obtained from blood samples drawn every 5 min (Fig. 2f). The fraction of inspired oxygen (FiO_2_) is varied to induce a range of SO_2_ levels, and the resulting paired measurements of HbO_2_/Hb concentrations and light intensities are used to calibrate the algorithm parameters (Analysis S1). The resulting FOP-derived HbO_2_/Hb concentrations and SO_2_ values show close agreement with BGA measurements throughout the calibration procedure (Fig. 2g and Fig. S8), confirming the validity of the calibrated model. This agreement is maintained across a physiologically relevant range of SO_2_ levels, demonstrating that the model is robust under varying physiological conditions. Furthermore, unlike the intermittent BGA sampling, the FOP provides continuous, real-time monitoring with a temporal resolution of 0.01 s.

To evaluate oximetric performance in a narrow-caliber vessel, we implant the FOP into the rabbit femoral artery^45,46^ (∼2 mm inner diameter) via the saphenous branch (Fig. S9), with a commercial pulse oximeter placed on a forepaw as a superficial reference. In a respiratory hypoxia model, reducing FiO_2_ from 100% to 10% causes SO_2_ to drop from 96.6% ±2.0% to a minimum of 60.9% ±15.1%, followed by a slight partial recovery due to homeostatic regulation, before returning to 96.0% ±3.0% upon reoxygenation (Figs. 2h and 2i). A milder protocol (20%→10%→20%) produces a smaller decline and faster stabilization (Fig. S10), consistent with a gentler physiological challenge^47^. The same optical signal also enables pulse-rate monitoring (Fig. 2h), which increases from 128 to 151 bpm during hypoxia and returns to baseline upon recovery, consistent with hypoxia-induced sympathetic activation^48^. While the superficial pulse oximeter shows similar overall trends during respiratory hypoxia, it records a greater decrease of SO_2_ and a longer equilibration time compared with the FOP. This indicates that peripheral tissues undergo more severe and prolonged desaturation during systemic hypoxia, consistent with a preferential redistribution of oxygen delivery to protect core organs^49^. These differences also highlight the FOP’s ability to perform oximetry within central vasculature that is inaccessible to superficial pulse oximetry.

In a femoral artery ligation model simulating intraoperative temporary occlusion^50^, we further assess the FOP’s capability to detect localized hemodynamic disturbances. The proximal femoral artery is temporarily occluded for 5 min to simulate surgical procedures such as temporary hemostasis or embolization, and distal SO_2_ is monitored continuously. Upon ligation, SO_2_ at the downstream site drops from 96.8% ± 1.5% to a minimum of 93.7% ± 1.6%, and remains suppressed throughout the occlusion period without signs of recovery. After releasing the ligature, SO_2_ rapidly recovers (Figs. 2j and 2k), indicating prompt restoration of oxygen delivery upon reperfusion. Meanwhile, the forepaw pulse oximeter shows no obvious SO_2_ change during ligation, confirming that the hypoxia was confined to the downstream territory of the occluded artery. This localized nature of the SO_2_ drop highlights a key limitation of systemic or superficial sensors in surgical settings, as they cannot detect region-specific ischemic events that occur deep within the operative field. By contrast, our FOP’s endovascular placement enables direct, real-time interrogation of the local hemodynamic status, which could facilitate the timely detection and management of focal complications during interventional procedures.

### FOP-based velocimetric sensing

The FOP system can also measure blood flow velocity via velocity-dependent dilution of a fluorescent dye (Fig. 3a). The decay signal of a dye bolus, which is injected through the microfluidic channel, is recorded by the fiber sensor through fluorescent photometry. Theoretical analyses (Fig. S11, Table S4 and Analysis S2) reveal that the fluorescence decay rate is proportional to local flow velocity, which can thereby be derived from the decay curve using a pre-established calibration model (Fig. S12). Compared with ultrasonic Doppler, thermodilution and conventional fluorescent photometry^51–54^, this approach offers highly localized interrogation, minimal dependence on operator experience and flow direction, and resistance to extravascular interference.

**Figure 3.**
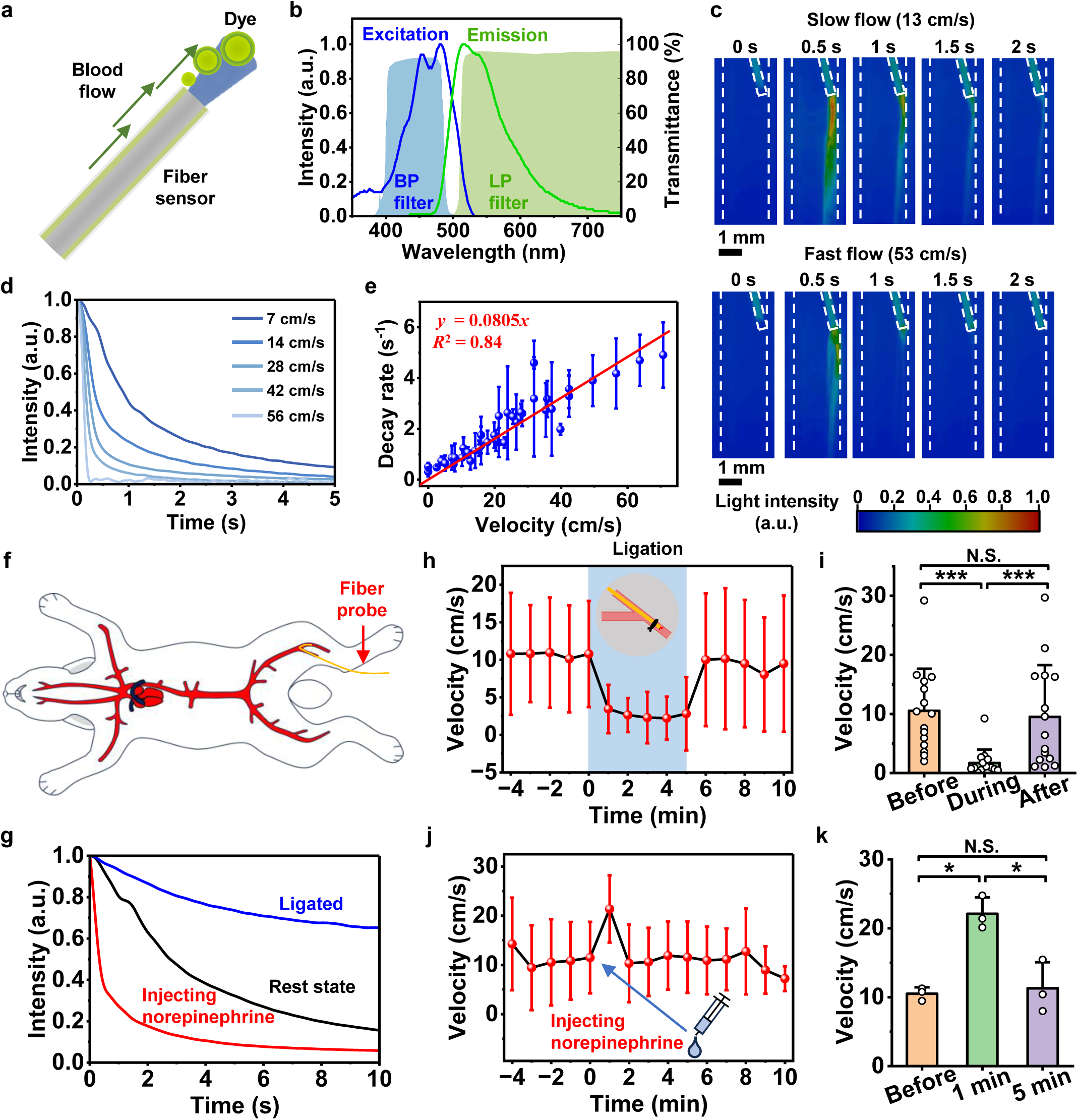
FOP-based blood velocimetry. **a.** Principle of blood flow velocity sensing. Fluorescent dye solution is injected through the microfluidic channel and diluted by the blood flow. The dilution induced fluorescence decay is detected by the FOP sensor and used to determine the blood flow velocity. **b.** Absorption and emission spectra of the fluorescent dye (sodium fluorescein), as well as transmittance spectra of bandpass (BP) and longpass (LP) filters used in velocity sensing. **c.** Time-lapse fluorescence images (pseudo-color) after dye injection into water with flow rates of 6 cm/s (upper) and 24 cm/s (lower). **d.** Fluorescence decay curves measured in blood flow of different velocities. **e**. Calibration curve showing the linear relationship between the inverse of the decay rate and the blood flow velocity. **f.** Illustration of in vivo blood velocity sensing experiment in a rabbit femoral artery model. **g.** Representative fluorescence decay curves measured within the artery in different conditions (rest state, ligation and injecting norepinephrine). **h.** Measured blood flow velocity change when acute ischemia and reperfusion are created by ligation in the proximal artery. **i.** Comparison of blood velocity results in the artery ligation model (15 trials, *n* = 8 rabbits). **j.** Measured blood velocity change upon norepinephrine injection. **k.** Comparison of blood flow velocity results in the norepinephrine injection model (3 trials, *n* = 3 rabbits). Paired two-tailed *t*-tests. \**p* < 0.05, \*\**p* < 0.01, \*\*\**p* < 0.001, N.S. *p* > 0.05. Data are presented as mean ± S.D.

We select sodium fluorescein as the signal indicator for its high quantum yield (>80%) and verified biocompatibility^55–57^, and set the working concentration at 0.1 mg/mL, below which its fluorescence remains proportional to concentration (Fig. S13), ensuring that the recorded decay curve accurately reflects the concentration decay. Its absorption and emission spectra are centered at ∼470 nm and ∼520 nm, respectively (Fig. 3b). Coupled to the FOB sensor, a photometric system comprises a 473 nm excitation laser, a photodetector, excitation/emission filters, and a programmable injection system for consistent injection volume and timing. Visualized in an artificial aqueous circulating system (Fig. 3c and Movie S4), fluorescence at the FOP tip rises rapidly after injection, and exhibits distinct decay curves between slow and fast flow velocities. Numerical simulations reproduce this behavior (Fig. S14 and Movie S5) and reveal a linear relationship between the decay rate and the flow velocity (Fig. S15), consistent with the theoretical analysis (Analysis S2). Simulations of probe implantation scenarios (Fig. S16) further indicate that FOP implantation causes only mild flow disturbance (velocity changes of 0.8%–6.2% across branches), lower than that induced by common catheterization (12%–50%)^58,59^.

Calibration in an artificial blood circulating system (Fig. S17a), with a commercial flowmeter as reference, shows steeper fluorescence decay at higher velocities (Fig. 3d) and a linear decay rate–velocity relationship (Fig. 3e), consistent with theoretical and numerical predictions (Fig. S11 and Fig. S15). Notably, data obtained in tubes of different diameters follow the same decay rate–velocity relationship (Fig. S17b), whereas the decay rate–flow rate relationship varies with tube size (Fig. S17c), confirming that velocity, rather than cross-sectional area or flow rate, governs the decay rate.

We then assess the FOP’s velocimetric performance in the rabbit femoral artery using the same implantation route as the oximetric experiments (Fig. 3f), examining hemodynamic conditions including resting state, vascular ligation, and pharmaceutical administration. Recorded decay curves (Fig. 3g) show that artery ligation slows the dilution process whereas norepinephrine injection accelerates it, demonstrating the FOP’s ability to resolve both decreases and increases in blood velocity. During ligation, blood velocity falls from 10.5 ± 6.9 cm/s to 1.7 ±2.2 cm/s, and recovers after release (Fig. 3h and 3i). The resting value agrees with reported values for the rabbit femoral artery^60^, and the stagnant flow during ligation confirms effective vascular occlusion. Comparing results in Figs. 2 and 3, the coordinated SO_2_ and velocity changes imply that impaired local SO_2_ regulation during occlusion arises from local ischemia; notably, the velocity reduction (∼84%) far exceeds the concurrent SO_2_ drop (∼3%), suggesting a modest oxygen supply–demand deficit compensated by increased oxygen extraction^61^. In the pharmacological model, intravenous norepinephrine, which simulates intraoperative hypotension treatment^62^, dramatically raises flow velocity within ∼1 min, recovering within ∼2 min (Figs. 3j and 3k). The resting velocity matched that of the ligation experiments. As an α-receptor agonist, norepinephrine induces vasoconstriction and elevates blood pressure^63^, and the observed femoral velocity increase confirms a systemic rather than local effect. Moreover, the rapid onset and short duration are consistent with its fast metabolism^64,65^.

### In vivo hemodynamic monitoring in a canine model

To demonstrate intraoperative monitoring in a clinically relevant setting, we monitor SO_2_ and blood velocity in a canine carotid artery model during balloon embolization (Fig. 4a). The canine carotid artery has an inner diameter of ∼2 mm and closely matches the target vasculature of deep interventional procedures in both caliber and access route^32,33,66,67^. Clinically, balloon embolization is routinely used for temporary flow control during stent and coil placement, yet carries a risk of complications^68,69^. Under DSA guidance, we implant a balloon catheter into the common carotid artery in a canine model, along with the FOP sensor positioned distal to the balloon (Fig. 4b and 4c, Fig. S18). Balloon inflation produces concurrent decreases in both SO_2_ and blood velocity. For oximetry, SO_2_ falls from 98.5% ± 1.6% to a minimum of 93.1% ± 3.2% upon inflation, and recovers to 96.2% ±3.4% after deflation (Figs. 4d and 4e). Meanwhile, blood velocity decreases from 23.5 ± 2.9 cm/s to a minimum of 5.6 ± 3.3 cm/s and returns to 14.3 ± 4.2 cm/s (Figs. 4f and 4g). Notably, peripheral SO_2_ recorded by a commercial multiparameter monitor remains unchanged throughout the procedure (Fig. 4d), whereas the intermittent velocity readings from a commercial ultrasonic Doppler velocimeter closely match the FOP measurement (Fig. 4f). These results demonstrate that the FOP is able to capture continuous, local hemodynamic changes in narrow vasculature that cannot be detected in real time by commercial non-invasive instruments during an interventional procedure.

**Figure 4.**
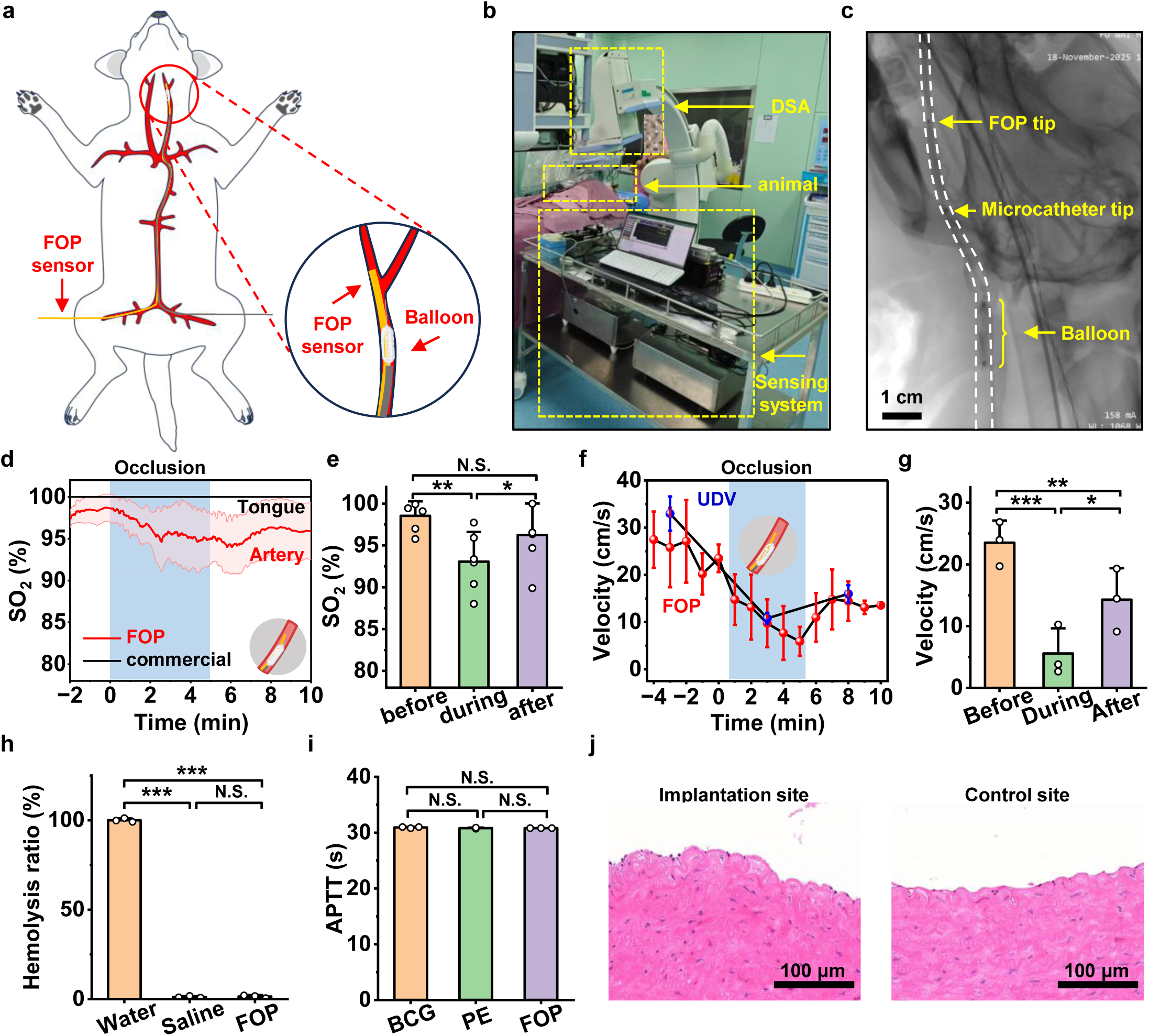
FOP-based blood oximetry and velocimetry during intraoperative vascular intervention in a canine model. **a.** Illustration of the intraoperative monitoring experiment. During the endovascular intervention, a balloon catheter is applied to embolize the carotid artery, while our FOP sensor is placed aside to monitor blood oxygenation and flow velocity. **b.** Photograph showing the intraoperative experiment, in which the catheter balloon and our FOP sensor are inserted under a surgical navigation system assisted by digital subtraction angiography (DSA). **c.** Angiographic image showing the insertion site. **d.** Measured SO_2_ change during balloon embolization. Red: results monitored by our FOP sensor within the artery; Black: results monitored by commercial surgical monitoring device externally attached to the tongue. **e.** Comparison of SO_2_ results in the balloon embolization model (4 trials, *n* = 2 dogs). **f.** Measured blood flow velocity change during balloon embolization. Red: results monitored by our FOP sensor within the artery; Blue: results measured by a commercial ultrasonic Doppler velocimeter (UDV) aside the common carotid artery. **g.** Comparison of flow velocity results in the balloon embolization model (3 trials, *n* = 2 dogs). **h.** Hemolysis rates tested for blood samples mixed in water, saline or our FOP sensor (*n* = 3 samples for each group). **i.** Activated partial thromboplastin times (APTT) measured for blood samples with different materials (*n* = 3 samples for each group). BCG: blank control group; PE: polyethylene. **j.** Hematoxylin and eosin (H&E) staining of vascular tissues in implantation and control sites. Paired two-tailed *t*-tests. \**p* < 0.05, \*\**p* < 0.01, \*\*\**p* < 0.001, N.S. *p* > 0.05. Data are presented as mean ± S.D.

Furthermore, these results illustrate the complementary value of dual-parameter sensing. They confirm that balloon embolization effectively restricted local blood flow while largely preserving oxygenation. Despite a significant reduction in velocity, the modest SO_2_ drop suggests a compensated oxygen supply, likely provided through residual flow from the contralateral carotid and vertebral arteries. Moreover, in contrast to the prompt flow rebound observed in the rabbit femoral artery, flow suppression here persisted beyond balloon deflation. This discrepancy likely reflects the distinct vascular beds and occlusion mechanisms involved across different ischemia–reperfusion models^70–72^, and it underscores the value of continuous local interrogation for capturing procedure-specific reperfusion dynamics^73,74^. Together, these findings demonstrate the FOP’s compatibility with microcatheter-based intervention and its potential for intraoperative monitoring in deep, narrow vasculature.

### Biocompatibility of the FOP sensor

As an acute implantable device in direct contact with blood, the FOP requires favorable hemocompatibility. We therefore assess three potential adverse responses: hemolysis, thrombosis, and arterial inflammation. First, the measured hemolytic rate of blood incubated with FOP segments is 0.25% ± 0.65%, statistically indistinguishable from the saline control (set as 0%; Fig. 4h and Fig. S19), indicating negligible hemolysis risk. Second, no significant differences in activated partial thromboplastin time (APTT) are observed among plasma incubated with FOP segments, polyethylene (PE), and a blank control (Fig. 4i), indicating that the FOP does not perturb the intrinsic coagulation pathway. Finally, histological analyses evaluate the interaction between the FOP tip and the vascular endothelium (Fig. 4j). The segments contacting the FOP exhibit a slightly ruffled, wavier endothelial morphology compared with uncontacted control parts, consistent with the mild mechanical effects expected for device intervention^75^. Neither segment shows dense inflammatory cell infiltration or aggregation of neutrophils, lymphocytes, or macrophages, indicating a quiescent inflammatory state. Collectively, these assays show low hemolysis, no significant change in APTT, and no overt inflammatory infiltration in the examined arterial sections during acute interventional monitoring.

## DISCUSSION

With a miniaturized structure, our FOP sensor operates within deep, narrow vasculature while providing continuous, localized monitoring of both blood oxygenation and flow velocity. This capability is enabled by the compact optofluidic probe design together with the oximetry and velocimetry principles developed for it, which yield measurement in close agreement with blood gas analysis and ultrasonic Doppler velocimetry, respectively. Notably, the value of the device lies not only in this agreement with reference standards, but in the local information it provides. Demonstrated in our rabbit and canine models, the FOP detects local hemodynamic behaviors in deep vasculature that are unable to be captured by the superficial sensor, and monitors distinct ischemia-reperfusion dynamics across different occlusion models.

Table S5 compares the FOP sensor with state-of-the-art endovascular hemodynamic sensors^20,22,27,76–80^. Compared with existing techniques, the FOP offers a unique combination of submillimeter size and dual-parameter sensing of blood oxygenation and flow velocity. The miniaturized size allows the FOP to enter deep, narrow vasculature for potential intraoperative monitoring, and the complementary oximetry and velocimetry provide a more comprehensive assessment of local hemodynamic status, potentially enabling earlier detection of procedure-induced ischemia and prolonged hypoperfusion.

Our target vessels (∼2 mm) are comparable in caliber to the proximal segments of the human anterior and middle cerebral arteries, and the access route mirrors clinical practice. Further translational implementations require validations in larger animals or non-human primates. While the current device is assembled from commercially available fiber and tubing components, advanced fabrication should yield probe structures with smaller footprints and comparable mechanical and optical performance, extending reach into deeper vasculature. Moreover, since the FOP can enter the cerebral circulation without damaging brain tissue, it may serve as a minimally invasive in situ sensor for continuous monitoring of cerebral metabolism.

Beyond hardware, sensing performance could be improved through data-driven signal processing. The current physics-based models are interpretable but may not exploit all information contained in the recorded spectra and decay information. Machine learning approaches, such as neural networks for spectral and temporal feature extraction, could improve measurement robustness, and accumulating intraoperative datasets may eventually enable early prediction of complications. Finally, the platform is not limited to oximetry and velocimetry. The microfluidic channel could be adapted for direct blood pressure measurement, and the fiber tip functionalized with biorecognition elements for fluorescence-based biochemical sensing. Given its demonstrated operation in deep, narrow vasculature, such a multifunctional device platform would be broadly applicable in interventional monitoring and biomedical research.

## Materials and Methods

### Fabrication of the FOP Sensor

The FOP sensor comprises a 250-μm-diameter PMMA optical fiber (ESKA CK10, Mitsubishi Chemical Group) and a polypropylene (PP) microfluidic tube (Qiujing Tubing Co., Ltd.) with an inner diameter of 400 μm and an outer diameter of 500 μm. In a typical fabrication procedure, the fiber was threaded into the tube, and both were fixed to a three-way connector (Figs. S1c and S1d). The connector’s inner channel has a diameter of 1 mm. The optical fiber and microfluidic tube enter through one end of the connector and were secured with epoxy adhesive (Scotch-Weld DP100 Plus, 3M). The side end of the tube was fixed to a 19-gauge blunt-tip syringe needle (outer diameter ∼1 mm). One opening of the microfluidic tube was embedded in the connector, forming a fluidic channel with the syringe needle. The optical fiber exits through the third end of the connector and was adapted with an FC ferrule for optical transmission. The connector ends were then filled with acrylic adhesive (5800, Gubaili) to render the tube watertight.

The tip of the optical fiber (Figs. S1a and S1b) was cropped so that it protrudes 0.3–0.5 mm beyond the microfluidic tube. A Pt-Ir ring (inner diameter 0.56 mm, outer diameter 0.61 mm, length 1 mm, Shenjie) was used as a radiopaque marker and fixed to the outer surface of the microfluidic tube at a distance of ∼5 mm from its tip.

### Extracorporeal Sensing System

The extracorporeal sensing system (Fig. 1i and Table S2) comprises two subsystems: optoelectronic and fluidic. The optoelectronic subsystem consists of a light source module, a photodetection module and a fiber-optic light path. Fiber-coupled LEDs (M530F3, M660FP1, and M810F2, Thorlabs, Inc.) and a fiber-coupled laser source (MBL-FN-473 nm, Changchun New Industries Optoelectronics Technology Co., Ltd.) served as light sources, driven and modulated by a lab-made current source. The current source outputs a time-stamp signal, indicating the switching time of every LED. The photodetection module consists of fiber-coupled Si-based photodetectors (LSSPD-1.2-2JKFC, Beijing Lightsensing Technology Co., Ltd.) for light signal collection, lab-made transimpedance amplifiers (20 kΩ) and an analog-to-digital converter (DAQ-M001, Suzhou Aumanyu Intelligent Technology Co., Ltd.) for signal conversion. A 2×1 fiber-optic coupler (core diameter 105 μm, 0.22 NA, Shenzhen Haichuang Optical Communication Co., Ltd.), 2 filters (480 nm short-pass and 510 nm long-pass) and fiber-optic patch cords established the optical path among the light source module, the photodetection module, and the FOP fiber.

The fluidic subsystem was based on a programmable injection pump (LSP001, Baoding Ditron Electronic Technology Co., Ltd.) equipped with a 1-mL glass syringe (Shanghai Gaoge Industrial and Trading Co., Ltd.) loaded with 0.1 mg/mL sodium fluorescein aqueous solution. The solution was prepared by mixing 10 mg sodium fluorescein, 5 mL of 625 U/mL sodium heparin solution, and 95 mL of normal saline. The syringe was connected to the FOP microfluidic channel via a Luer lock. In a typical velocimetric trial, the pump injected a 10 μL bolus of dye solution at a flow rate of 130 mm/min, followed by a second 10 μL bolus at 60 mm/min after 2 s.

### Data processing

For oximetry (Fig. S6), the intensity of each wavelength was first separated from the time-stamp signal. The signal was then filtered with a zero-phase Butterworth low-pass filter (*f*_pass_ = 48 Hz, *f*_stop_ = 50 Hz, *R*_pass_ = 5 dB, *R*_stop_ = 30 dB) and decomposed by PCHIP-based empirical mode decomposition (EMD). The signal was reconstructed from the residual and intrinsic mode functions with frequency < 0.3 Hz or > 0.1 Hz, then transformed by a Daubechies4-based stationary wavelet transform (SWT) at level 4 and reconstructed from the stationary wavelet approximation coefficients. The pulse signal was extracted by peak detection, from which the pulse rate was calculated. The signal was further filtered with a zero-phase Butterworth low-pass filter (*f*_pass_ = 1 Hz, *f*_stop_ = 1.5 Hz, *R*_pass_ = 5 dB, *R*_stop_ = 30 dB). Blood oxygen saturation was then calculated from the filtered signal according to the algorithm described in Analysis S1.

For velocimetry (Fig. S12), the maximum light intensity was identified and recorded for each dye-injection trial. The intensity decay curve was numerically differentiated (Δ*t* = 2 ms), and the minimum derivative min(d*I*/d*t*) was recorded. The decay rate was obtained as abs[min(d*I*/d*t*)]/max(*I*). Blood velocity was calculated from this decay rate using the calibration relationship described in Analysis S2.

### Mechanical characterization

Mechanical properties were characterized for the optical fiber, the microfluidic tube and the integrated FOP. The strain–stress curves were obtained from a tensile test machine (Ez test, Shimadzu Corporation), from which elastic modulus and elastic limit were calculated. 10-cm-length samples were tested with a tensile rate of 10 mm/min. Bending stiffness was measured by self-weight tests. In a typical test, one end of the fiber or tube was fixed to the edge of a metallic frame, allowing the free end to hang under its own weight, and the beam deflection was measured. Samples of different lengths were tested to establish a length–deflection relationship. Based on the beam deflection formula *δ* = *qL*^4^/(8*EI*) (where *δ* is deflection, *L* is length, *E* is elastic modulus, and *I* is moment of inertia), the bending stiffness (*EI*) was calculated.

### Optical characterization

The wavelength-dependent transmission loss of the optical fiber was measured using the optoelectronic subsystem. Fiber samples of different lengths (1, 2, 5, and 10 m) were prepared, terminated with FC ferrules, and connected between the light sources and photodetector. The output power of the light sources was maintained at a constant level, and the transmitted intensity was recorded by the data acquisition module. From these measurements, a length–intensity relationship was obtained. Based on the light transmission formula *I* = *I*_0_*e*^−*aL*^ (where *I* is the transmitted intensity, *I*_0_ is the incident intensity, *a* is the attenuation coefficient, and *L* is the fiber length), the transmission loss of the fiber was calculated.

Bending stability was evaluated using a 1 m optical fiber. Bends of different angles (0°, 90°, 180°, 270°and 360°) were formed by wrapping the optical fiber around a 5-mm-diameter PTFE rod. Light of different wavelengths was provided by the light source module, and the transmitted intensity was recorded for each bending angle by the photodetection module.

### Sensing principle and validation

#### Optical simulation

Optical simulation was performed with TracePro 7.3 (Lambda Research Corporation, trial version). An artery-in-tissue model (Fig. S7a) was constructed, consisting of a cylindrical blood-filled artery (inner diameter 1.08 mm, outer diameter 1.6 mm) surrounded by a rectangular gray matter block with a side length of 2 cm, and an overall model length of 6 cm. Two sets of optical parameters were used, corresponding to blood at 98% and 0% SO_2_ (Table S3). A disc light source (250 μm diameter, 22° divergence angle) was placed at the center of the artery to simulate light emission from the optical fiber, and a detector disc of the same size was placed 1 μm behind the source to simulate light collection. A total of 500,000 rays were traced.

The energy received by the detector disc was recorded as an indicator of the light intensity collected by the optical fiber. To obtain the spatial distribution of light in blood, a surface perpendicular to the radial direction of the disc was defined as the measurement plane, and its power distribution was extracted for further analysis.

#### Fluidic simulation

Fluidic simulation was performed with Ansys Fluent 2021 R1 (ANSYS, Inc.). Two models were constructed: an FOP-in-vessel model to simulate the dilution procedure at different flow velocities, and a branching artery model to evaluate how endovascular implantation affects the local blood flow field. Table S4 shows the fluidic parameters of blood and dye aqueous solution used in simulation.

In the FOP-in-vessel model (Fig. S14a), a sensor consisting of a fiber (diameter 0.25 mm, length 1 mm) and a tube (inner diameter 0.4 mm, outer diameter 0.5 mm, length 0.5 mm) was embedded in a cylindrical blood domain (diameter 2 mm, length 10 mm). The annular space between the fiber and the tube was set as the dye inlet, with its velocity determined by the flow rate of the dye solution delivered by the FOP. The annular space between the tube and the cylindrical wall was set as the blood inlet, with its velocity varied across simulations to represent different flow conditions. The opposite opening of the cylinder was set as a pressure outlet with zero gauge pressure. Transient simulation was performed with 2,500 time steps of 2 ms each, and the dye concentration distribution on the longitudinal section of the cylinder was recorded.

In the branching artery model (Fig. S16a), the main artery (diameter 4.2 mm) bifurcates into two branches of 3 mm and 2 mm diameter, with a total artery length of 20 mm. Steady-state simulation was used. The main artery opening was set as a velocity inlet (100 cm/s), and the two branch openings were set as pressure outlets with zero-gauge pressure. Ten cross-sections along the branching arteries were selected as measurement planes, and the average velocity at each plane was recorded after simulation. Three conditions were simulated: the original state without the implanted FOP, the FOP (diameter 0.5 mm) positioned in the larger branch, and the FOP positioned in the smaller branch.

#### Ex vivo porcine blood test

A yeast solution was prepared by mixing 1 g of yeast with 10 mL of normal saline and incubating it in a 37 °C water bath for 10 min. Porcine whole blood (9 mL) treated with sodium citrate anticoagulant (Shanxi Junma Biotechnology Co., Ltd.) was placed in a centrifuge tube and incubated in a 37 °C water bath for 10 min. Yeast solution (1 mL) was then added to the blood and mixed thoroughly, after which the mixture was sealed with 1 mL of soybean oil and returned to the water bath. The FOP sensor was immersed in the mixture, and the returned light intensity was recorded by the optoelectronic subsystem for 60 min.

#### Artificial circulation system tests

An artificial circulating system was constructed to validate the fluorescent dilution method (Fig. S17a). The system comprised a liquid reservoir (Deerkang Medical Co., Ltd.), two brushless pumps (Nanjing Xinke Co., Ltd.), an electromagnetic flowmeter (AK66, Yihai Flowmetry Co., Ltd.), a hemostasis valve (Zhongchao Plastic Co., Ltd.), a PTFE flow control valve, and PTFE tubing connecting them into a closed loop. The pumps provided constant driving force, the flow control valve regulated the flow velocity, and the flowmeter provided standard measurement. The hemostasis valve allowed FOP insertion without liquid leakage.

Normal saline and porcine blood (Shanxi Junma Biotechnology Co., Ltd.) were used as circulating media in fluorescence microscopy and in vitro calibration, respectively. For fluorescence microscopy, the dilution process around the sensor tip was recorded at 13 cm/s and 53 cm/s with a lab-made fluorescence microscope equipped with a 470 nm LED excitation source and a CCD camera. For calibration, different flow velocities were established by adjusting the flow control valve, and the tube downstream of the hemostasis valve was interchanged among tubes of different inner diameters to investigate whether cross-sectional area affects the fluorescence dilution curve.

### In vivo experiments

#### Animals

All animal procedures were reviewed and approved by the Institutional Animal Care and Use Committee (IACUC) of Tsinghua University and the IACUC of Fuwai Hospital, under approved protocol numbers 18-SX1.G22 and 18-SX1.G25 (Tsinghua University) and 0108-3-6-HX(Y)-62 (Fuwai Hospital). All in vivo procedures were conducted in strict accordance with institutional and national guidelines for animal experimentation.

Adult New Zealand rabbits (3–4 kg) were purchased from Laboratory Animal Research Center of Tsinghua University and housed individually under standard conditions at 18–22 °C on a 12 h light/dark cycle with access to food and water ad libitum.

Adult beagle dogs (10–15 kg) were purchased from Animal Experimental Center of Fuwai Hospital, Chinese Academy of Medical Sciences and Peking Union Medical College and housed individually under standard conditions at 18–25 °C on a 12 h light/dark cycle with access to food and water ad libitum.

#### In vivo oximetry calibration

New Zealand rabbits were anesthetized by intramuscular administration of a Zoletil–Xylazine combined anesthetic (5 mg/mL Zoletil 50, 10 mg/mL xylazine, 1.3 mL/kg). The animal was placed in the supine position, and the inguinal region was shaved and aseptically prepared. A longitudinal skin incision (approximately 2 cm) was made directly over the femoral artery. Blunt dissection was performed to expose the femoral artery, which was isolated over a length of approximately 1 cm. A 20G indwelling catheter was inserted into the femoral artery and advanced toward the abdominal aorta. The catheter was secured within the vessel with a ligature, and a hemostasis valve connected to the catheter hub allowed the FOP sensor to be introduced without blood leakage. The FOP sensor, sterilized with 75% alcohol and heparinized with heparin saline in advance, was inserted through the catheter, and gently advanced retrogradely into the abdominal aorta. The position of the FOP tip was confirmed by measuring the insertion depth of the FOP advanced into the artery.

A blood gas analyzer (BGA, ABL90 FLEX, Radiometer Medical ApS) was used as the calibration standard. The contralateral femoral artery was exposed, and a 26G indwelling catheter was inserted, gently advanced retrogradely to the abdominal aorta, secured to the skin, and flushed with heparinized saline. Every 5 min, a blood sample (approximately 0.5 mL) was withdrawn into a pre-heparinized blood gas syringe and immediately analyzed. A facial mask was fixed over the nose and mouth of the rabbit and connected to a gas mixer of O_2_ (high-purity oxygen, Linde Gas) and N_2_ (high-purity nitrogen, Linde Gas). The flow rates of O_2_ and N_2_ were adjusted to control the fraction of inspired oxygen (FiO_2_). The FOP sensor and BGA measured blood oxygen saturation simultaneously throughout the procedure. After calibration, the FOP sensor and indwelling catheters were withdrawn, both femoral arteries were tied off permanently. While the rabbit remained under deep anesthesia, it was transferred to a sealed chamber for euthanasia by nitrogen asphyxiation. Nitrogen gas was introduced into the chamber at a flow rate of 1 L/min, gradually displacing ambient air. Respiration and heartbeat were continuously monitored, and death was confirmed by the irreversible cessation of respiratory movements and heartbeat for at least 5 minutes.

#### Femoral artery implantation via the saphenous artery

Following the same anesthesia, animal positioning, and aseptic preparation described for in vivo calibration, a longitudinal skin incision (approximately 2 cm) was made over the femoral triangle. The femoral artery was exposed by blunt dissection, and the saphenous artery, a branch of the femoral artery, was identified and isolated over a length of approximately 1 cm. Two 4-0 nylon sutures were passed beneath the isolated segment. The distal end of the saphenous artery was permanently ligated, and a loose temporary ligature was placed around the proximal portion. A small arteriotomy was created just proximal to the distal ligature using microsurgical scissors. The sterilized and heparinized FOP sensor was inserted through the arteriotomy and gently advanced into the femoral artery, then delivered downstream to the intended location (1–2 cm distal to the branching point of saphenous artery). The position of the FOP tip was confirmed by the light spot at its optical fiber.

After the FOP sensor was properly positioned, the corresponding experimental protocol (respiratory hypoxia, artery ligation, or pharmacological model) was performed as described in the following subsections. After all measurements, the FOP sensor was withdrawn, the proximal saphenous ligature was tied off permanently, and the animal was euthanized.

#### Respiratory hypoxia model

Following FOP implantation in the femoral artery as described above, a veterinary pulse oximeter probe was clipped to a forelimb, and heart rate and oxygen saturation were continuously monitored throughout the procedure. A facial mask was fixed over the nose and mouth of the rabbit, and the fraction of FiO_2_ was varied. In one set of experiments, FiO_2_ was set to 1000%, 10%, and 1000% sequentially, with each stage lasting 5 min. In another set, the three 5 min stages were set as 20%, 10%, and 20%. The FOP sensor continuously measured blood oxygen saturation throughout these procedures.

#### Artery ligation model

Following FOP implantation in the femoral artery as described above, a veterinary pulse oximeter probe was clipped to a forelimb, and heart rate and oxygen saturation were continuously monitored. A nylon suture was looped around the proximal femoral artery and tightened to completely occlude blood flow. Occlusion was maintained for 5 min, after which the ligature was released to allow reperfusion. The FOP sensor continuously measured both blood oxygen saturation and blood velocity during this procedure, beginning 5 min before injection.

#### Pharmacological model

Following FOP implantation in the femoral artery, the rabbit ear was shaved and aseptically prepared. A 26G indwelling catheter was inserted into the marginal ear vein and secured to the skin. Norepinephrine solution was prepared in advance by diluting 0.5 mL of 0.1% norepinephrine in normal saline with 19.5 mL of normal saline. Norepinephrine solution was then injected through the catheter at a dose of 0.042 mg/kg over approximately 30 s. Immediately after administration, the catheter was flushed with 5 mL of heparinized saline to ensure complete delivery. The FOP sensor continuously measured blood velocity during this procedure, beginning 5 min before injection.

#### Canine vascular intervention experiments

Beagles were premedicated with propofol and anesthetized with inhaled isoflurane, endotracheally intubated, and mechanically ventilated throughout the procedure to maintain normal respiration. A multiparameter monitor continuously recorded heart rate, blood pressure, oxygen saturation, and end-tidal CO_2_. Both inguinal regions were shaved and aseptically prepared, and a longitudinal skin incision (approximately 3–4 cm) was made over the femoral triangle on each side. Blunt dissection was performed to expose the femoral artery, which was isolated over a length of approximately 1.5–2 cm. Proximal and distal vascular control was achieved with vessel loops. An 18G puncture needle was introduced into the isolated segment of one femoral artery. The needle was then withdrawn and a 5F arterial sheath was placed and secured. The same procedure was performed on the other femoral artery to establish bilateral vascular access. Systemic heparinization was then initiated with a bolus of heparin (100 U/kg) and maintained with supplemental doses as needed.

Under digital subtraction angiography (DSA; Innova 2010-IQ, GE Healthcare), a 3F guiding catheter was then introduced through the left femoral sheath and navigated into a common carotid artery over a 0.018-inch guidewire. The wire was removed, and the FOP sensor—heparinized with heparin saline and disinfected with povidone-iodine in advance—was advanced coaxially through the guiding catheter to the target segment under roadmap guidance (the common carotid artery in one beagle and the internal carotid artery in the other). Correct positioning was confirmed by Angiography. Subsequently, via the right femoral sheath, a second guiding catheter and a balloon microcatheter were advanced into the same common carotid artery, with the balloon tip positioned proximal to the FOP tip.

After placement, the balloon was inflated with contrast medium. Occlusion was maintained for 5 min, after which the balloon was deflated by aspirating the inflation fluid. The FOP sensor continuously measured blood oxygen saturation and flow velocity during the procedure, beginning 5 min before balloon inflation. An ultrasonic Doppler velocimetry system (Vivid E9, GE Healthcare) simultaneously measured blood velocity in the common carotid artery.

After all measurements, all catheters, the FOP, and sheaths were removed. Both femoral arteries were ligated at the puncture sites to ensure hemostasis, and the inguinal skin incisions were closed in layers with absorbable sutures. The animals were returned to their cages for recovery. Postoperative analgesia and antibiotics were administered according to the institutional protocol, and the animals were monitored daily.

### Biocompatibility characterization

#### Hemolysis rate tests

Diluted porcine blood was prepared by mixing 8 mL of anticoagulated porcine whole blood (Shanxi Junma Biotechnology Co., Ltd.) with 10 mL normal saline. Three groups were tested, each in triplicate: an experimental group (normal saline containing FOP segments), a negative control (normal saline alone), and a positive control (deionized water). For the experimental group, 1 cm FOP segments were placed in centrifuge tubes, and 10 mL of normal saline was added to each tube. For the negative and positive controls, 10 mL normal saline or deionized water was added, respectively.

To evaluate the hemolysis rate, 0.2 mL of diluted porcine blood was added to each tube. After thorough mixing, the tubes were incubated for 60 min in a 37 °C water bath. The liquid in each tube was then collected and centrifuged for 5 min at 2500 rpm. The supernatant was transferred to a well plate, and the absorbance at 545 nm was measured using a plate reader (Varioskan Lux 3, Thermo Fisher Scientific). The hemolysis rate was calculated according to the following equation^79^:

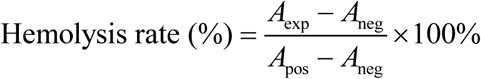

where *A*_exp_ is the absorbance of the experimental group, *A*_neg_ is the absorbance of the negative control, and *A*_pos_ is the absorbance of the positive control.

#### APTT tests

Bovine plasma samples were purchased from Shanxi Junma Biotechnology Co., Ltd. Three groups were prepared, each in triplicate. In the two experimental groups, a 1 cm FOP segment or a 1 cm ×1 cm polyethylene (PE) piece was added to 15 mL of plasma and shaken at 60 rpm at 37 °C for 15 min. The materials were then removed, and the plasma was retained for APTT testing. For the blank control group (BCG), plasma was shaken without foreign objects.

APTT measurements were performed on a RAC-1830 coagulation analyzer (Rayto Life and Analytical Sciences Co., Ltd.). The three plasma groups were placed in the sample rack, and the analyzer automatically performed the following procedure. 50 μL of plasma and 50 μL of APTT reagent were aspirated and incubated at 37 °C for 3 min. 50 μL of 0.025 mol/L CaCl_2_, pre-warmed to 37 °C, was then added, and the time from CaCl_2_ addition to clot formation was recorded as the APTT result.

#### Histology evaluation

After euthanasia of the beagles, proximal and distal segments of the common carotid arteries (approximately 1 cm each) were harvested. The proximal segments had been in contact with the microcatheter, balloon catheter, and FOP sensor, whereas the distal segments had not. The samples were rinsed with PBS and fixed in 4% paraformaldehyde at 4 °C for 24 h. After fixation, the tissues were dehydrated through a graded ethanol series, cleared in xylene, and embedded in paraffin. Serial sections of 4–5 μm thickness were cut alongside the axial direction using a rotary microtome, mounted on glass slides, and dried overnight at 37 °C. The sections were deparaffinized in xylene, rehydrated through a descending ethanol series, and stained with hematoxylin and eosin (H&E). After staining, the sections were dehydrated, cleared in xylene, and coverslipped with neutral mounting medium. Stained sections were examined under a light microscope, and representative images were captured.

## Acknowledgements

This work is supported by the National Natural Science Foundation of China (NSFC) (T2425003 and 52272277, to X.S.; 32571534 to L.Y.; 62422501 to H.D.), the National Key Research and Development Program of China (2024YFB4607704 to X. L.), Beijing Natural Science Foundation (Z220015 and L254019 to L.Y., L2606020 to H.D.), Noncommunicable Chronic Diseases-National Science and Technology Major Project (2024ZD0525805, to H.D.), Tsinghua-Toyota Joint Research Fund (20253930040 to X.S.). The authors acknowledge characterization work supported by Center of Intelligent Sensing and Precise Testing Technology in Optics and Photonics, BIT and Tsinghua Nanofabrication Technology Center.

## Author contributions

X. S., X. L. and L. Y. developed the concepts. B. Z., Z. Y., D. L., D. S. and X. C. performed device and circuit design, fabrication and characterization. B. Z. and X. W. performed the simulations. B. Z., X. L., Z. Y., X. W., W. K., H. Z., D. L., J. C., Z. X., Y. W. and Q. Y. designed and performed biological experiments. X. L., H. D., L. Y. and X. S. provided tools and supervised the research. B. Z. and X. S. wrote the paper in consultation with other authors.

## Data and materials availability

All data needed to evaluate the conclusions in the paper are present in the paper and/or the Supplementary information.

---

**Figure S1.**
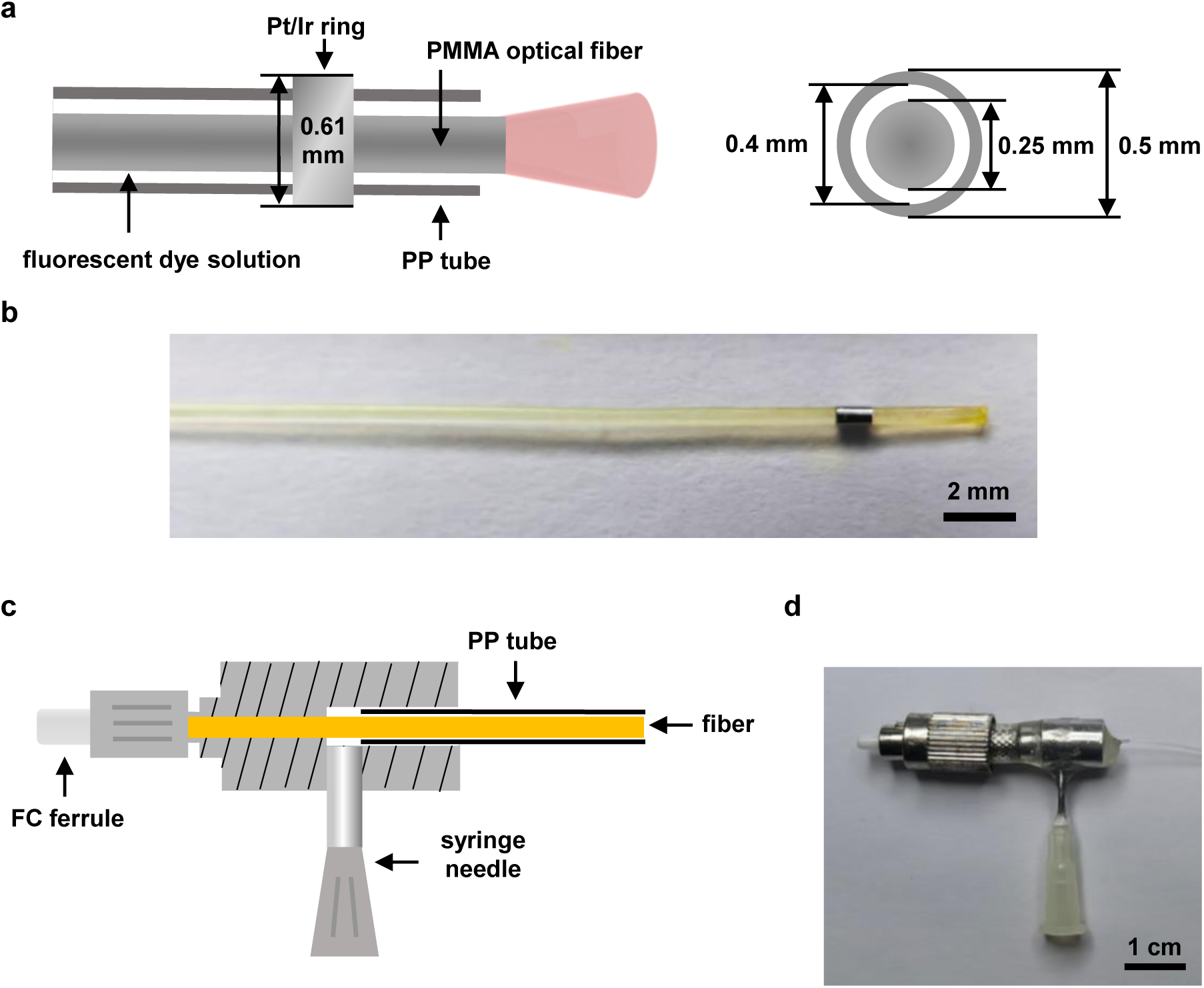
Structure layout of the FOP sensor. **a.** Tip of the probe, including a PMMA optical fiber for light emission and reception, a PP tube as a microfluidic channel for dye injection, and a metallic (Pt/Ir) ring as a radiopaque marker. Left: side view; Right: cross-sectional view. **b.** Photograph of the FOP sensor tip**. c.** Connection part of the probe, linked with optical system and microfluidic system. **d.** Photo of the connection part.

**Figure S2.**
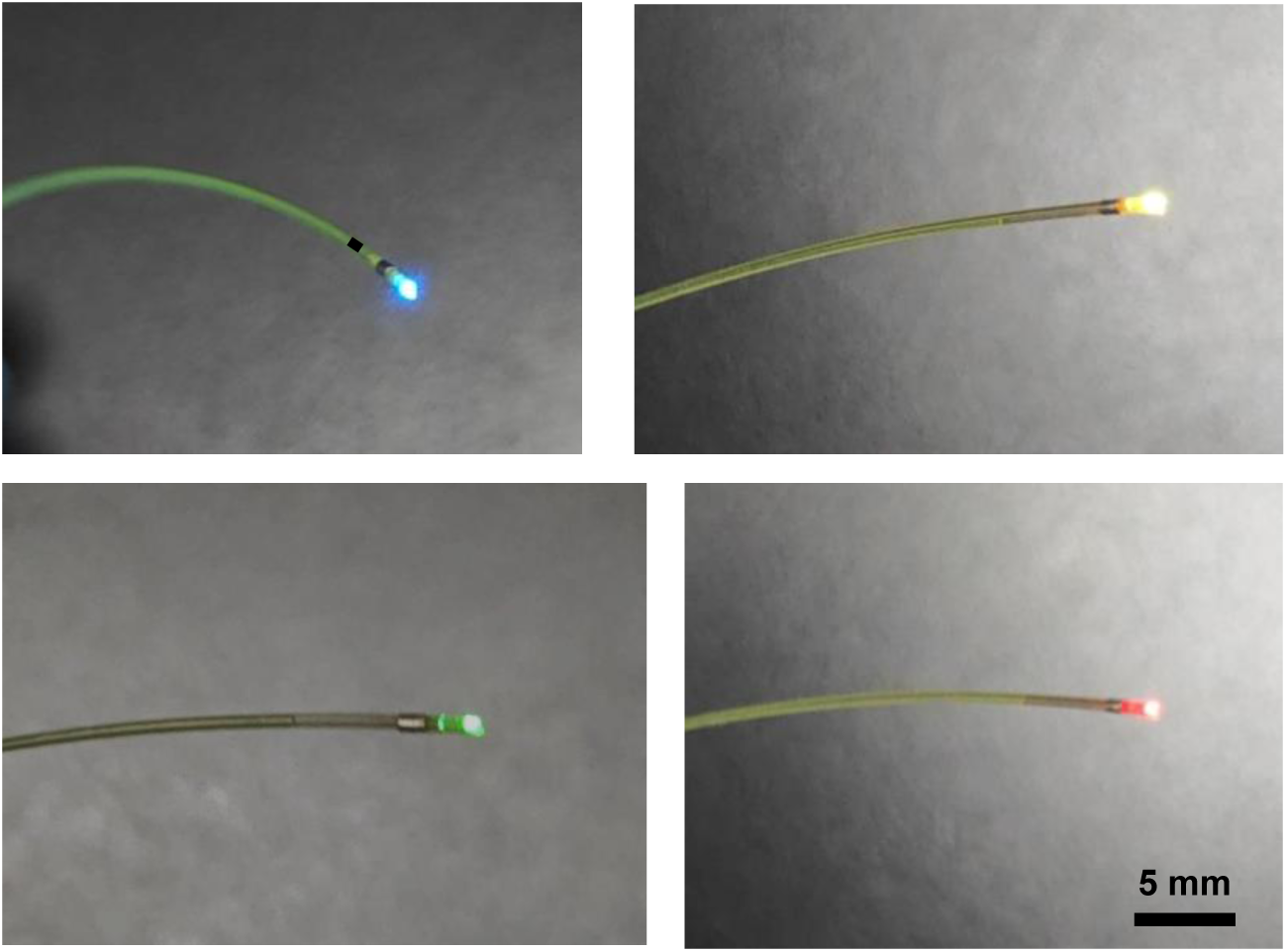
Photographs of an FOP sensor emitting light of different wavelengths.

**Figure S3.**
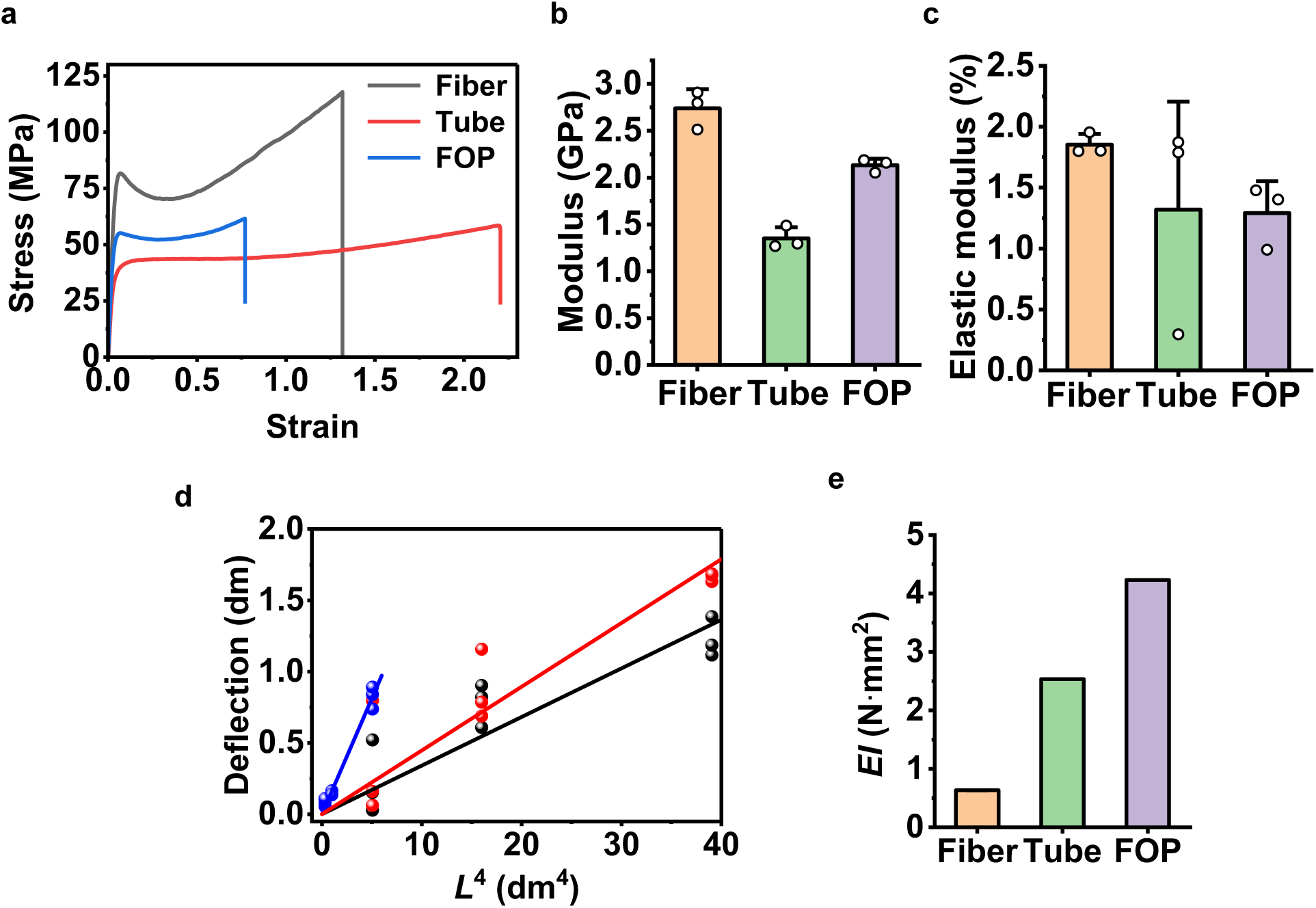
Mechanical characterization of the PMMA fiber, the PP tube and the integrated FOP sensor. **a.** Strain–stress curves. **b.** Elastic modulus. **c.** Elastic limit. **d. S**elf-weight beam deflection tests. **e.** Bending stiffness.

**Figure S4.**
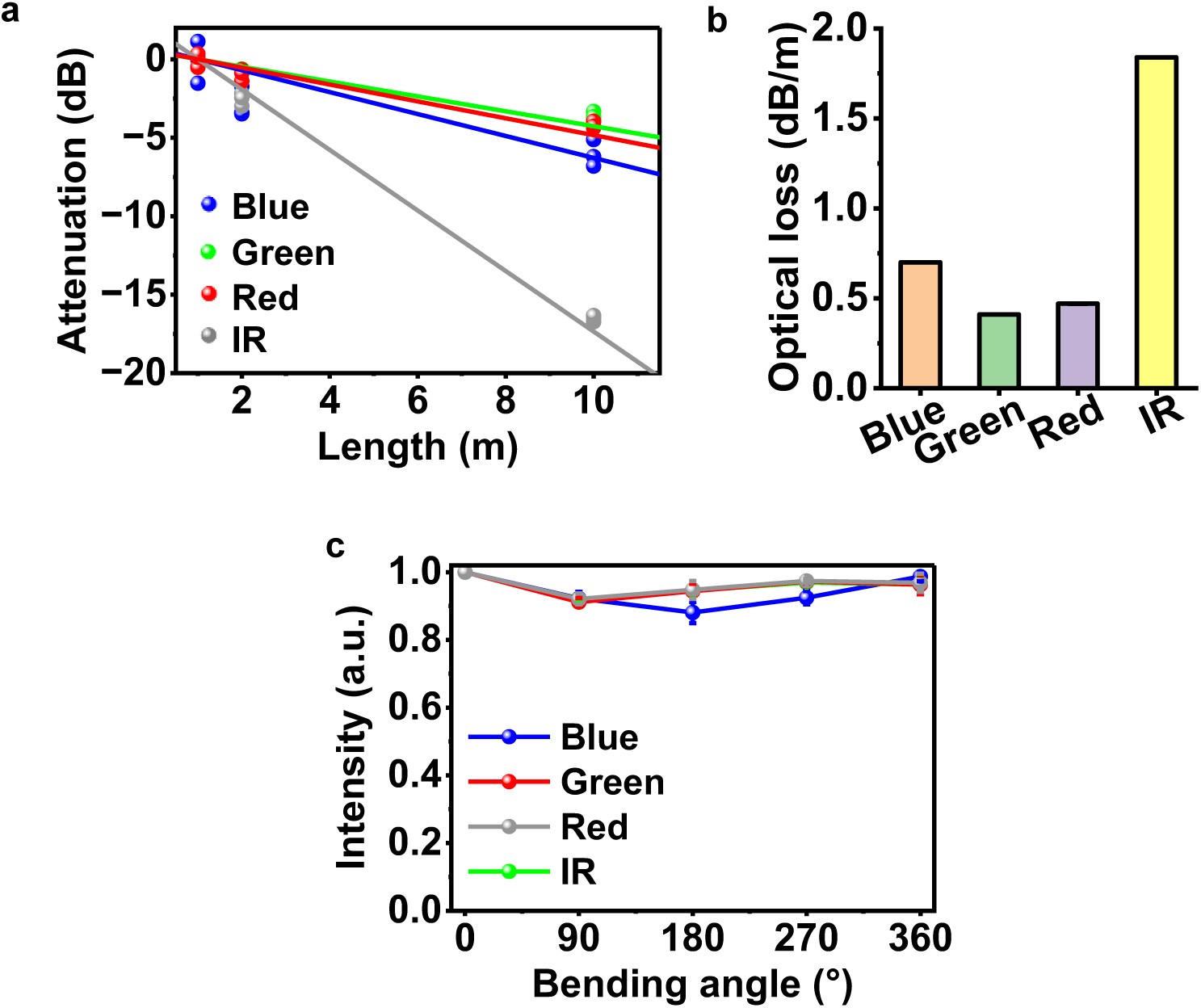
Optical characterization of the PMMA optical fiber in the FOP sensor. **a.** Attenuation measured through optical fiber of different lengths, at different wavelengths (blue: 470 nm; green: 530 nm; red: 660 nm; IR: 810 nm). **b.** Transmission loss at different wavelengths. **c.** Light intensity through optical fiber with different bending angles (bending radius: 5 cm).

**Figure S5.**
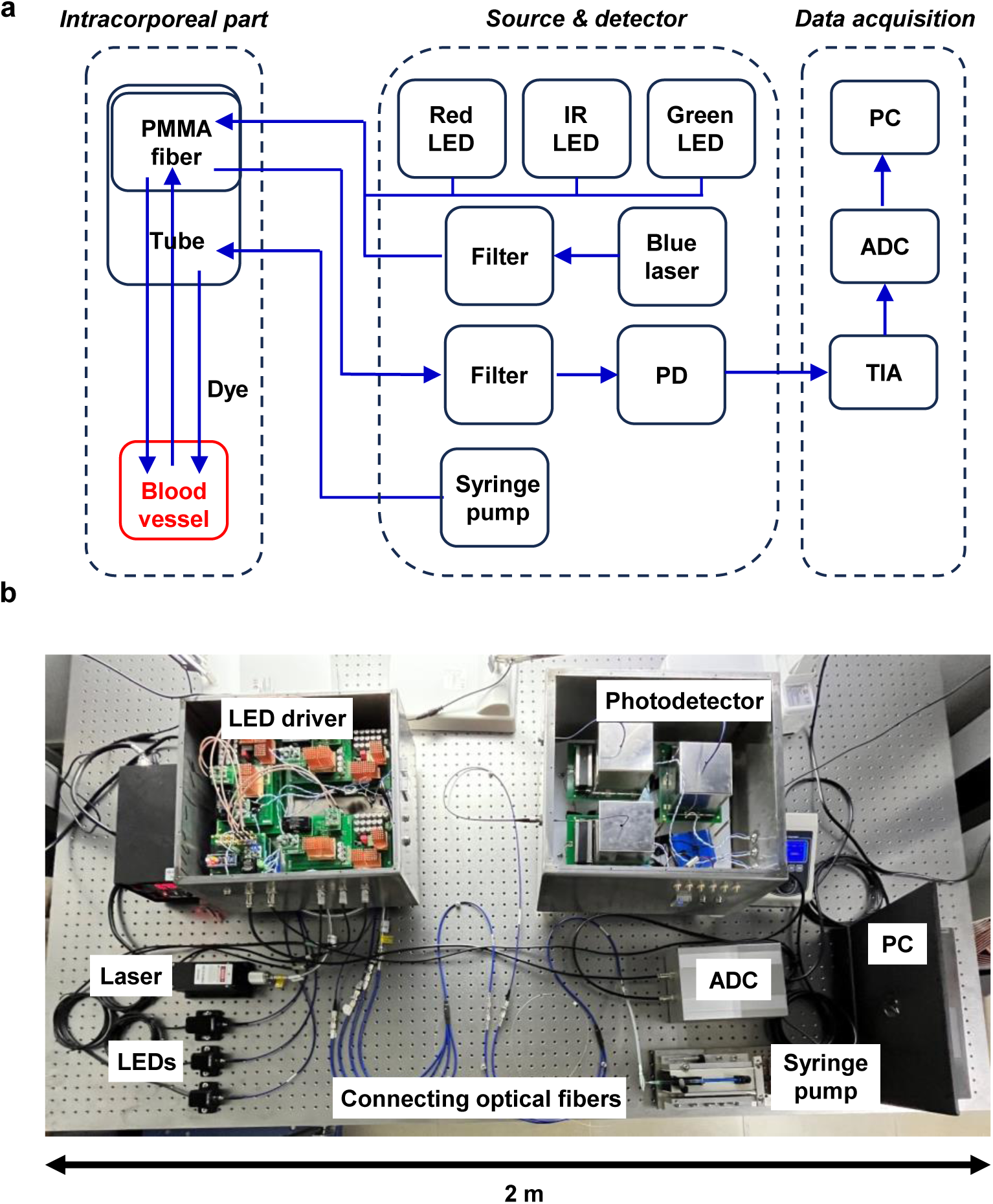
FOP sensing system for blood oximetry and velocimetry. **a.** Block diagram of the entire system, which comprises 3 sub-systems: the intracorporeal part for light delivery and dye injection, the active device part (light sources, detectors and syringe pump), the data acquisition and processing unit. **b.** Photograph of the whole system.

**Figure S6.**
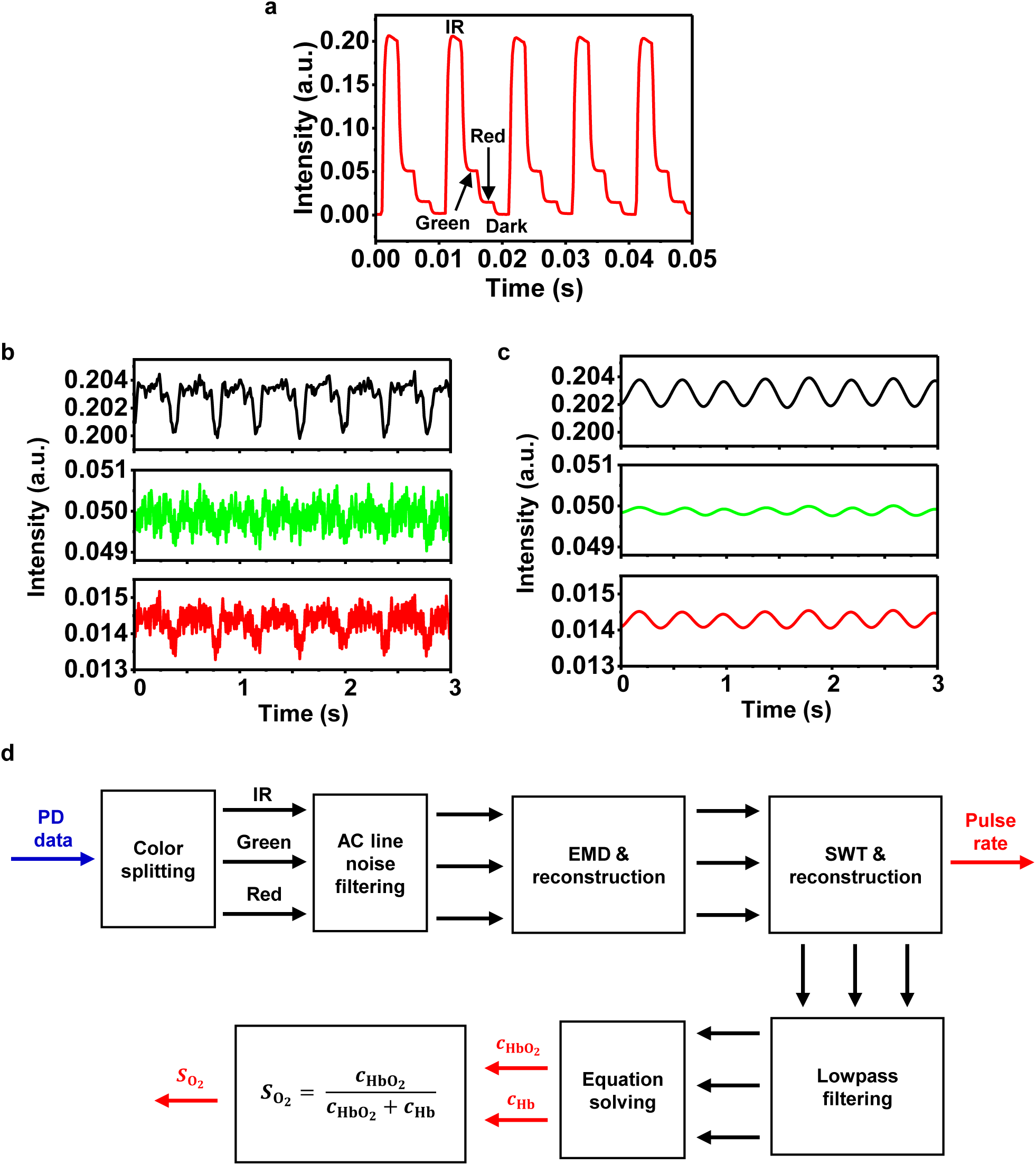
Signal processing procedure for blood oxygenation calculation. **a.** Typical raw light intensity data recorded by our FOP sensing system. Light of different wavelengths (810 nm, 530 nm and 660 nm) are emitted and collected alternately. **b.** Typical light intensity data of different colors after raw data splitting. **c.** Typical pulsed data after filtering. **d.** Block diagram of data processing procedure. EMD: empirical mode decomposition, SWT: stationary wavelet transformation.

**Figure S7.**
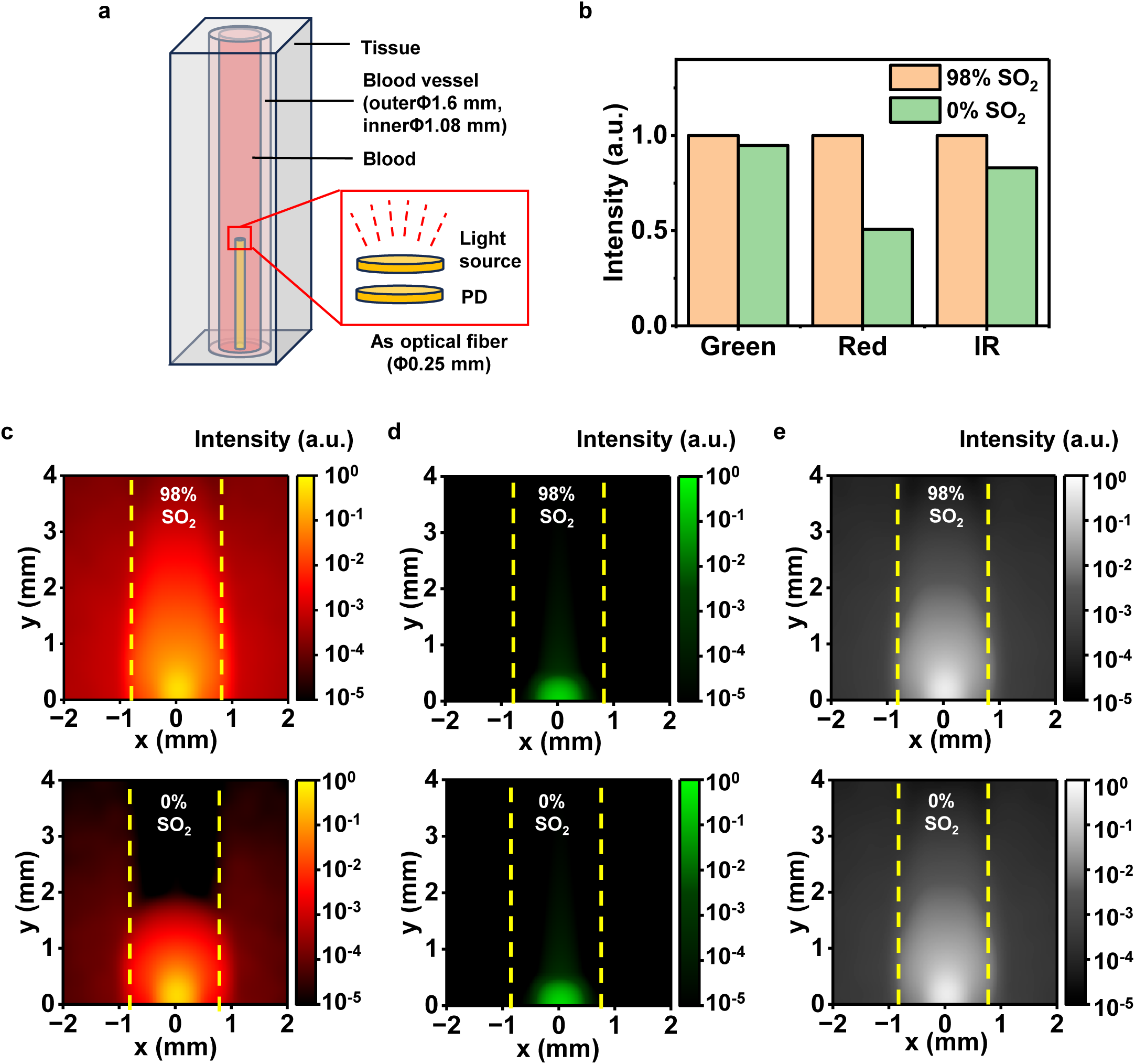
Numerical Simulation of optical intensity distribution in the vascular tissue with different oxygenations. **a.** Optical model used in the simulation, consisting of blood, vessel wall, rounding tissue, a light source, and a photodetector. **b.** Received light intensity at different wavelengths for blood oxygenation of 98% SO_2_ and 0% SO_2_. **c–e.** Simulated intensity distribution at 660 nm **(c)**, 530 nm **(d)**, and 810 nm **(e)** for blood with 98% SO_2_ (upper) and 0% SO_2_ (lower).

**Figure S8.**
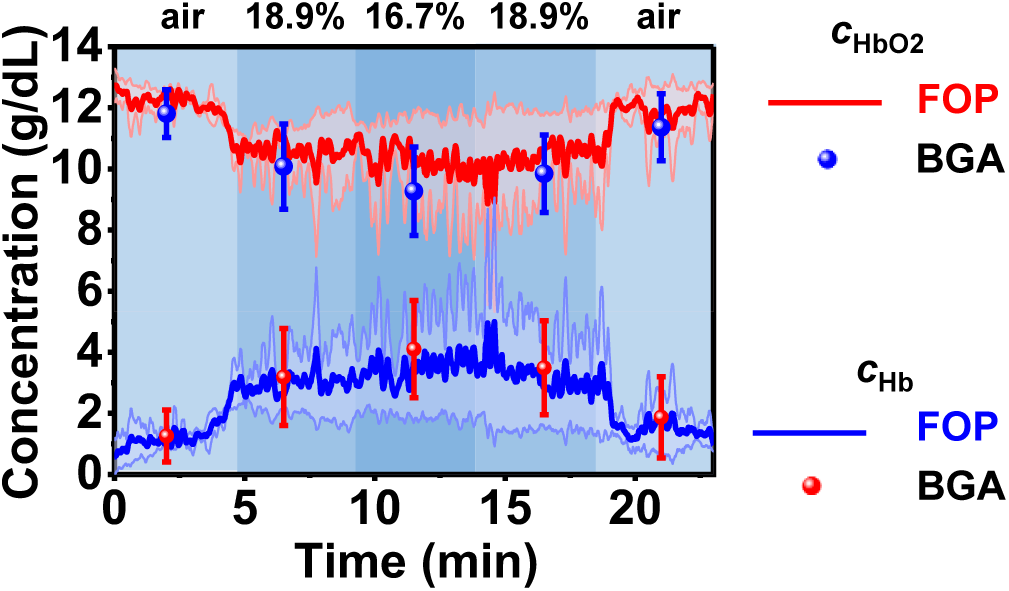
In vivo measured concentrations of hemoglobin. Concentrations of oxyhemoglobin (*c*_HbO2_) and deoxyhemoglobin (*c*_Hb_) measured at different fractions of inspired O_2_ (*F*_iO2_) by our FOP sensor (line), compared with data simultaneously collected by a commercial blood gas analyzer (BGA) (dot) (2 trials, *n* = 2 rabbits). These data are used to determine *S*_O2_ results in Fig.2g.

**Figure S9.**
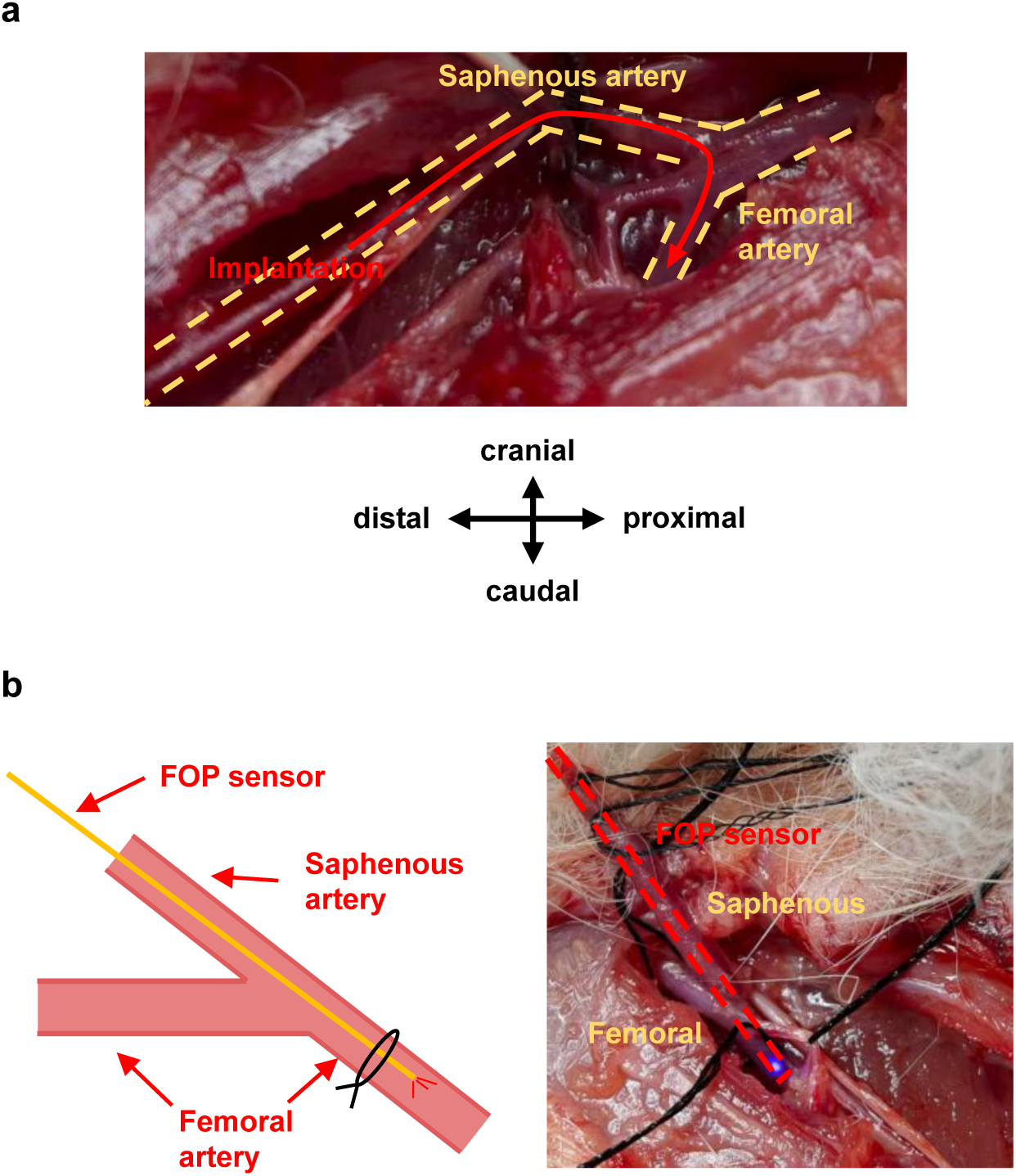
Surgical setup of probe insertion in rabbit femoral artery. **a.** location of implantation: proximal end of saphenous artery. **b.** implantation route. Saphenous artery is opened to form an entry, through which the FOP sensor is inserted into the distal part of the femoral artery.

**Figure S10.**
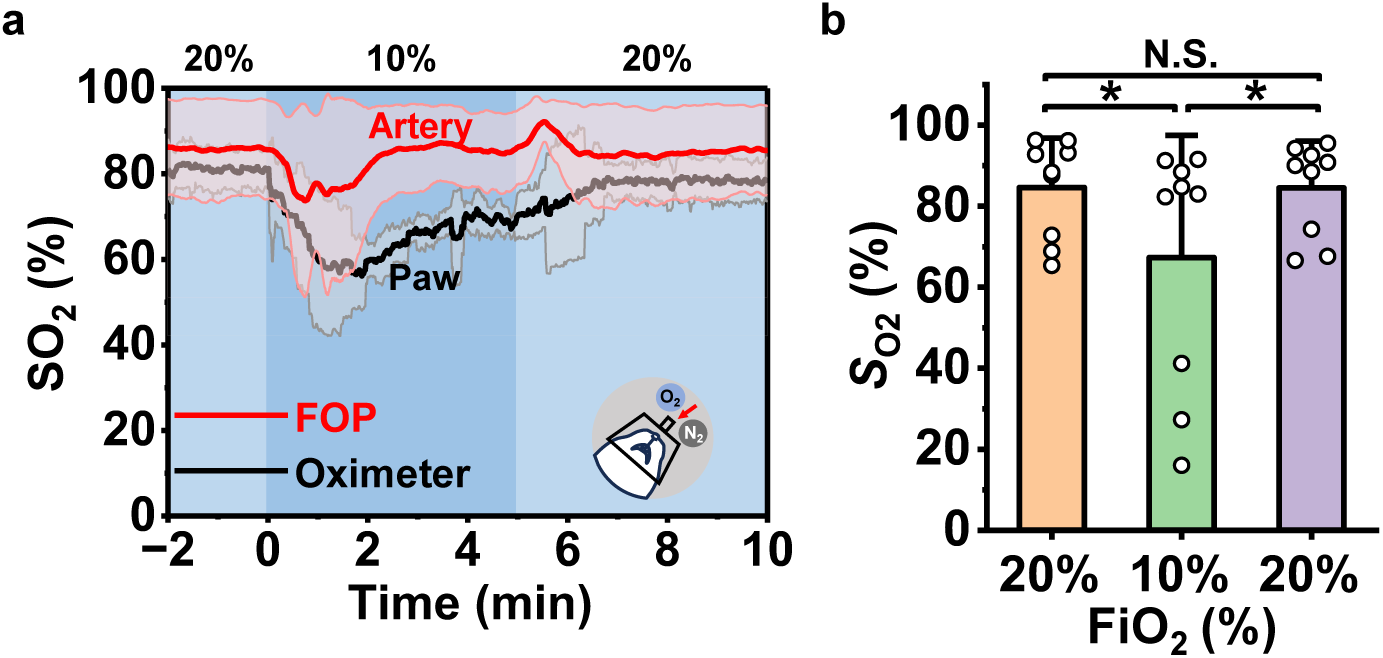
In vivo oximetric results in another FiO_2_ changing pattern. **(a)** SO_2_ results measured at different FiO_2_ (20%–10%–20%). Red: results detected by our FOP sensor within the femoral artery; Black: results collected externally by a commercial pulse oximeter mounted at the paw. **(b)** Comparison of SO_2_ results at FiO_2_ varying from 20% to 10% and to 20% (9 trials, *n* = 3 rabbits). Paired two-tailed *t*-tests. \**p* < 0.05, \*\**p* < 0.01, \*\*\**p* < 0.001, N.S. *p* > 0.05. Data are presented as mean ± S.D.

**Figure S11.**
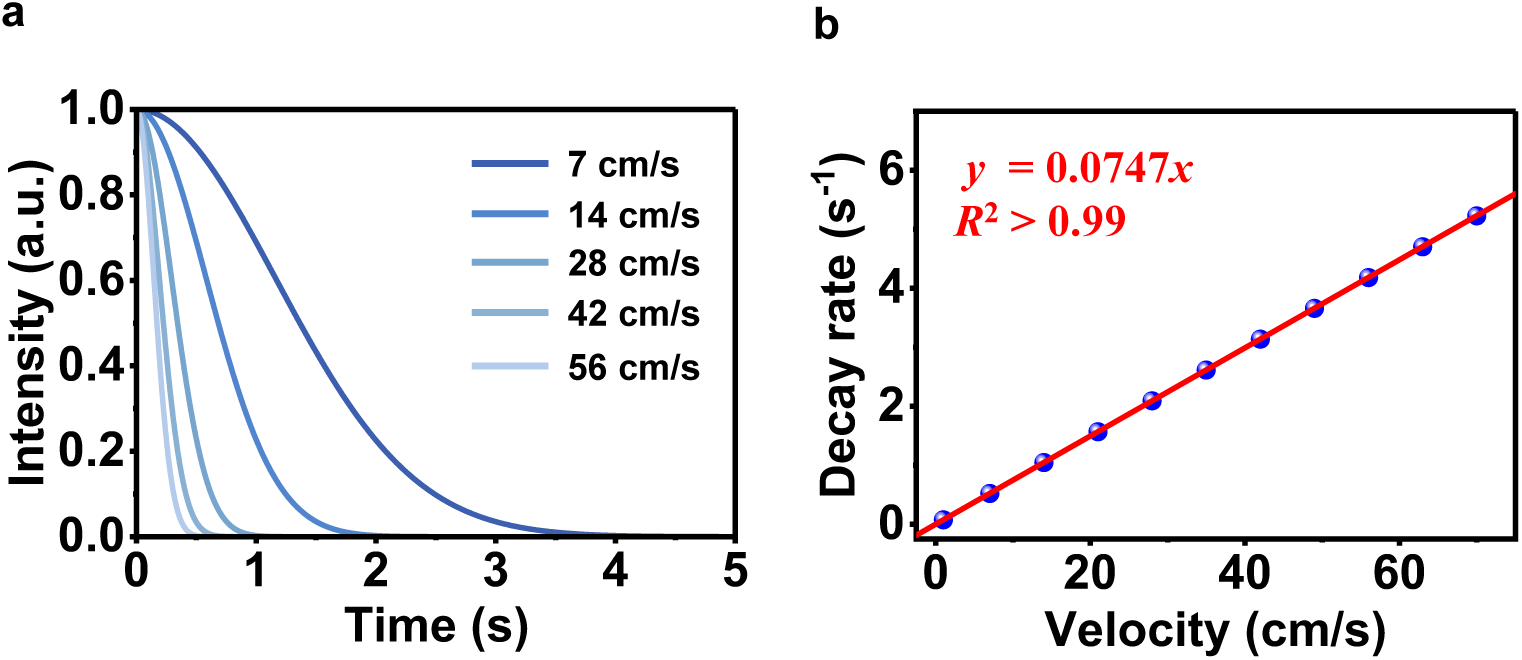
Theoretical relationship between fluorescent dilution procedure and flow velocity based on Analysis S2. (a) Time-lapsed fluorescence intensity decay. (b) Corresponding relationship between decay rate and blood velocity.

**Figure S12.**
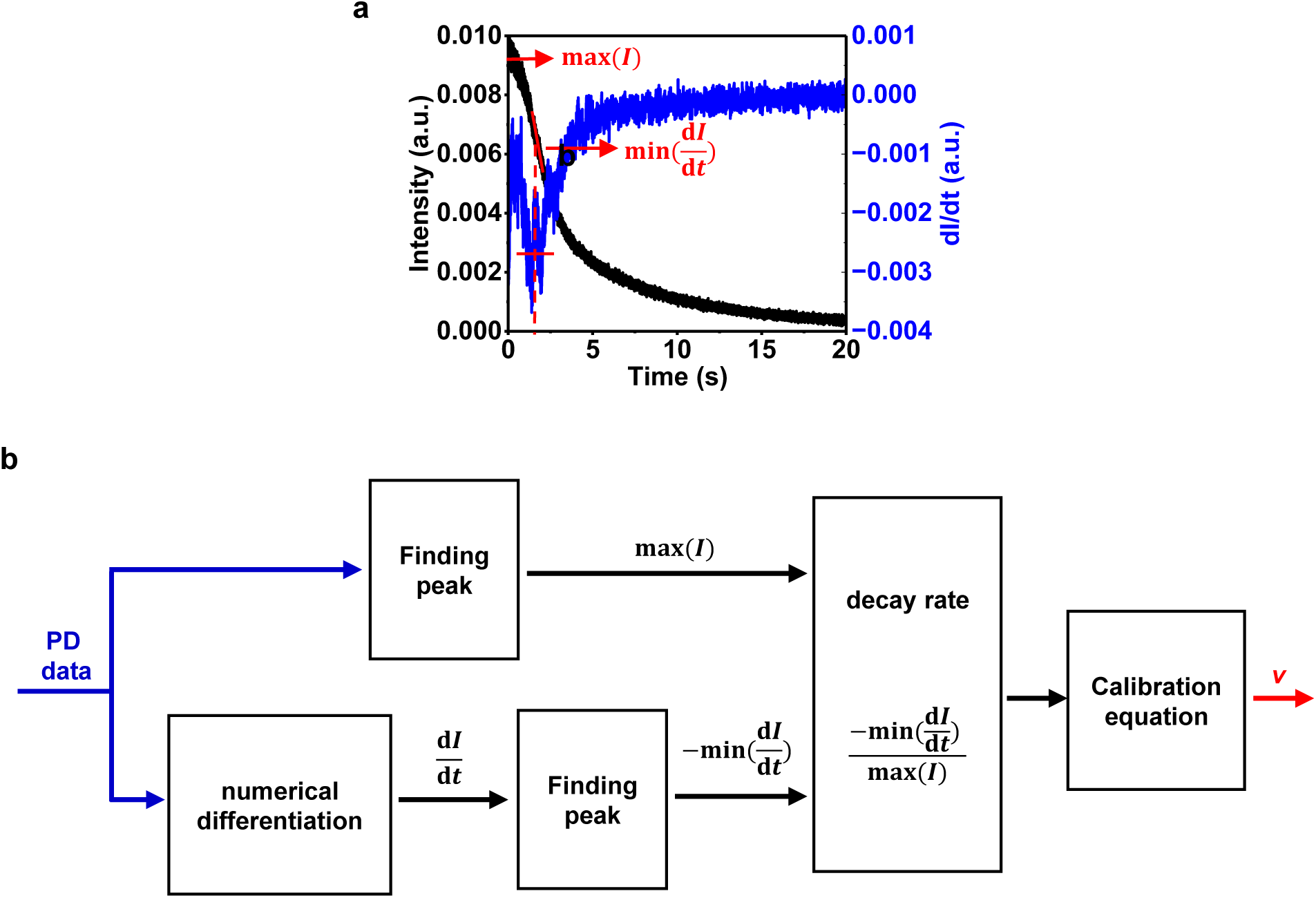
Signal processing procedure for blood velocity calculation. **a.** Typical feature extracted from a measured fluorescence decay curve. **b.** Block diagram of data processing procedure.

**Figure S13.**
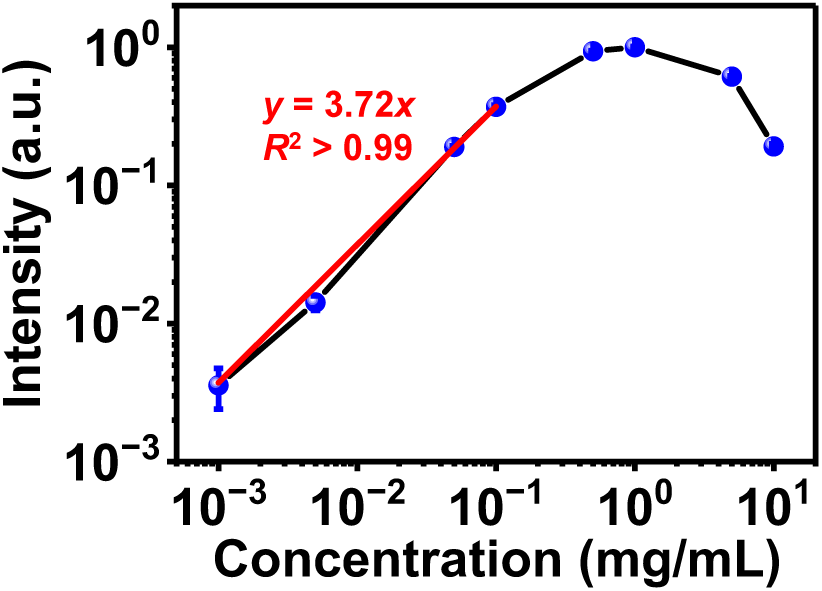
Measured photoluminescence (PL) intensity as a function of the dye (sodium fluorescein) concentration. Data are recorded by immersing an FOP sensor in aqueous solution of different sodium fluorescein concentrations. Red line shows linear fitting result for concentrations below 0.1 mg/mL.

**Figure S14.**
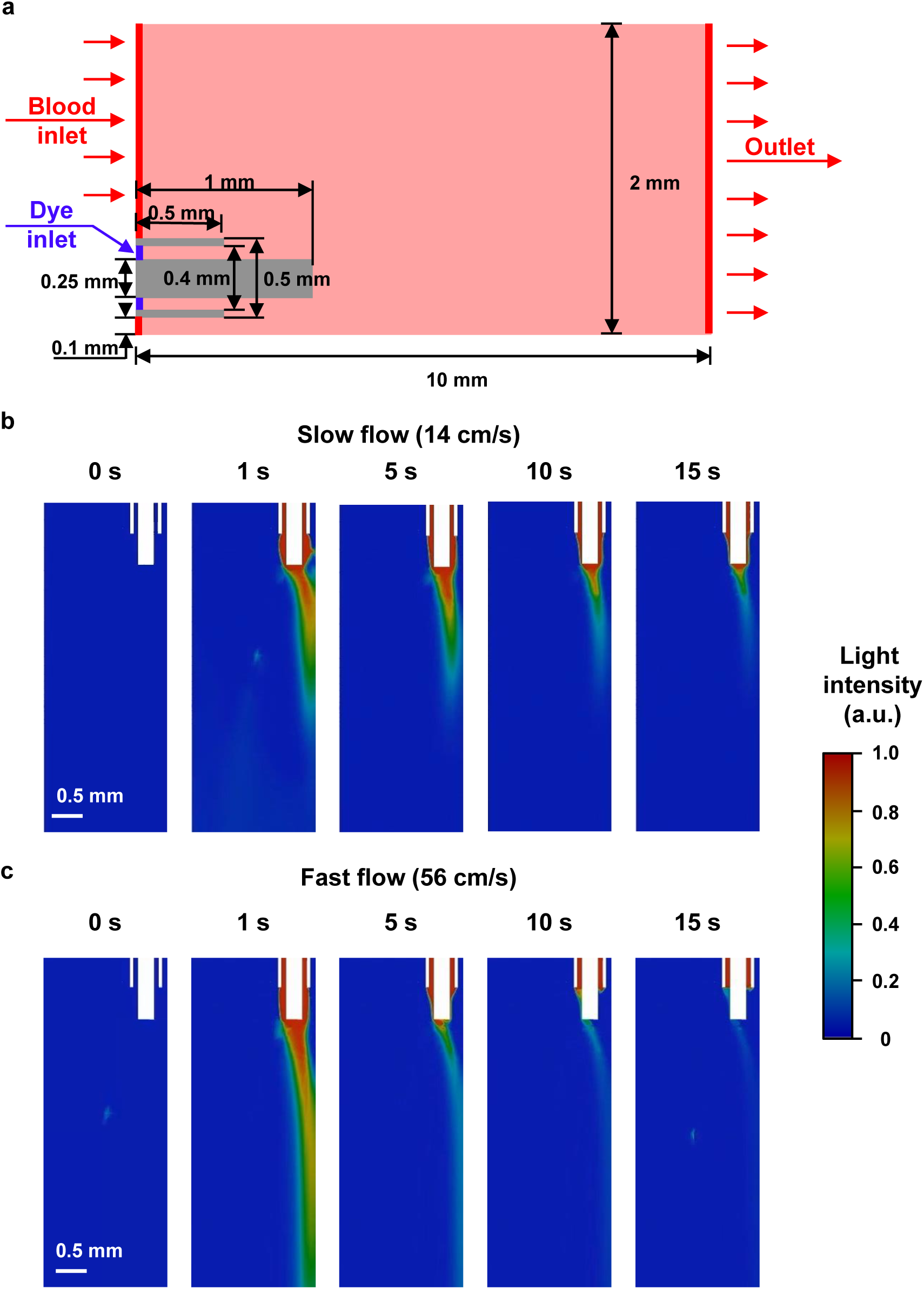
Numerical simulation of the fluorescent dilution procedures. (a) Fluidic model used in numerical simulation, consisting of a 2-mm-diameter blood vessel and the FOP. (b,c) Time-lapse images of the dye dilution process in blood flows of different velocities. Two fluidic velocities are used, (b) 14 cm/s and (c) 56 cm/s.

**Figure S15.**
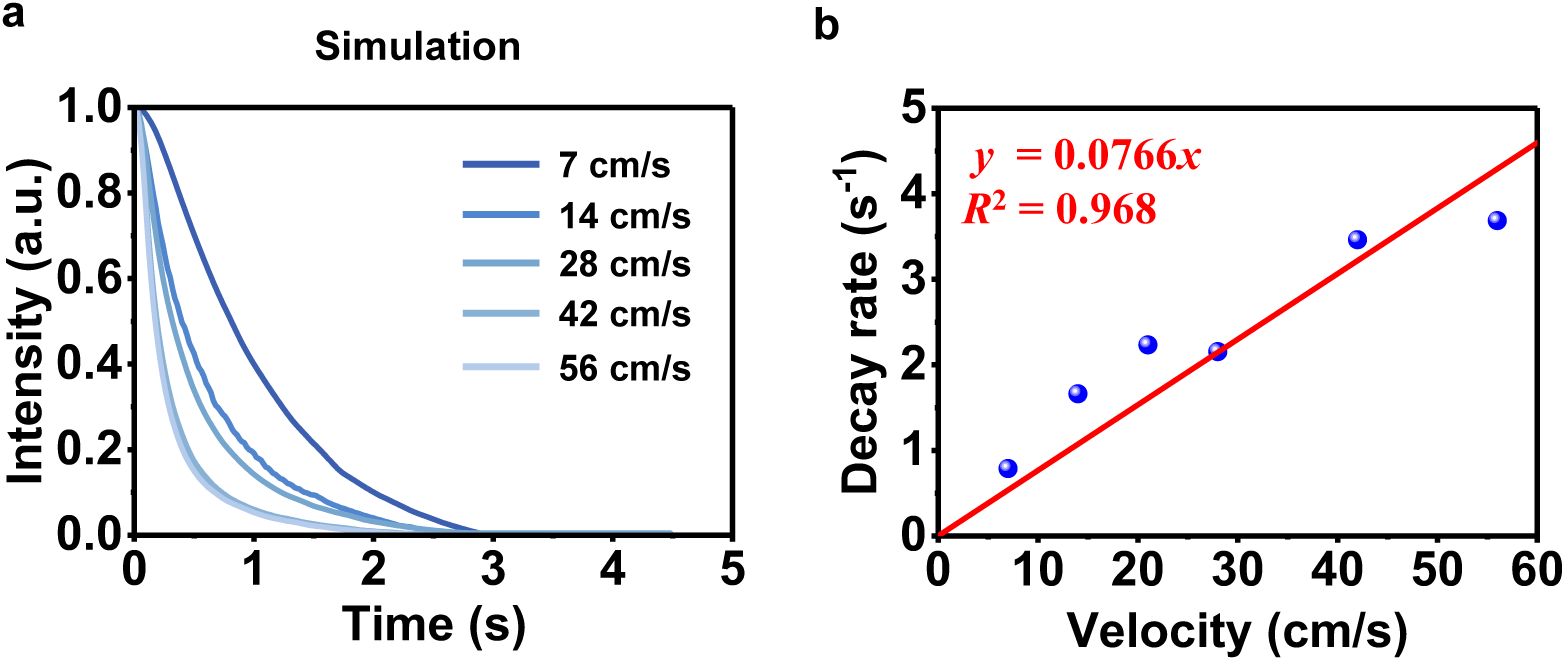
Numerical simulation results on the relationship between fluorescent dilution procedure and flow velocity. (a) Time-lapsed fluorescence intensity decay. (b) Corresponding relationship between decay rate and blood velocity.

**Figure S16.**
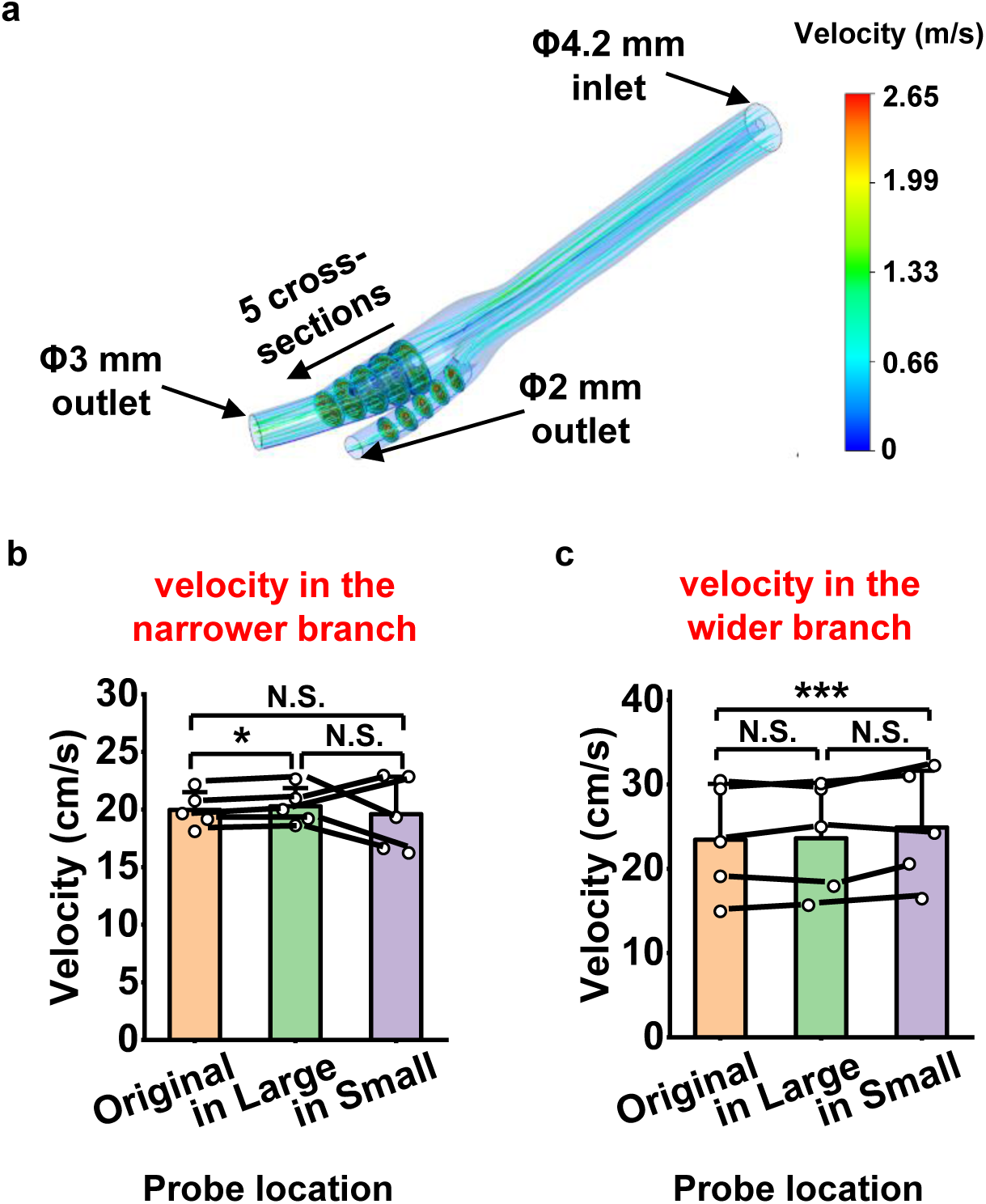
Hydrodynamic simulation to study the influence on blood velocity induced by inserting the FOB sensor. **a.** model layout. In this model, a bifurcating artery structure is studied, in which the sensor is placed in one of the two branches. **b, c.** statistical analysis of the velocity change in the narrower branch **(b)** and the wider branch **(c)**. Paired two-tailed *t*-tests (*n* = 5 cross-sections for each group). Data are presented as mean ± S.D. Every dot in the graph corresponds to average velocity of a cross-section along the blood vessel. For the vessel where fibrous sensor was placed, polarity of velocity change varies across different cross-sections. While for the other vessel, the velocity increases for all cross-sections.

**Figure S17.**
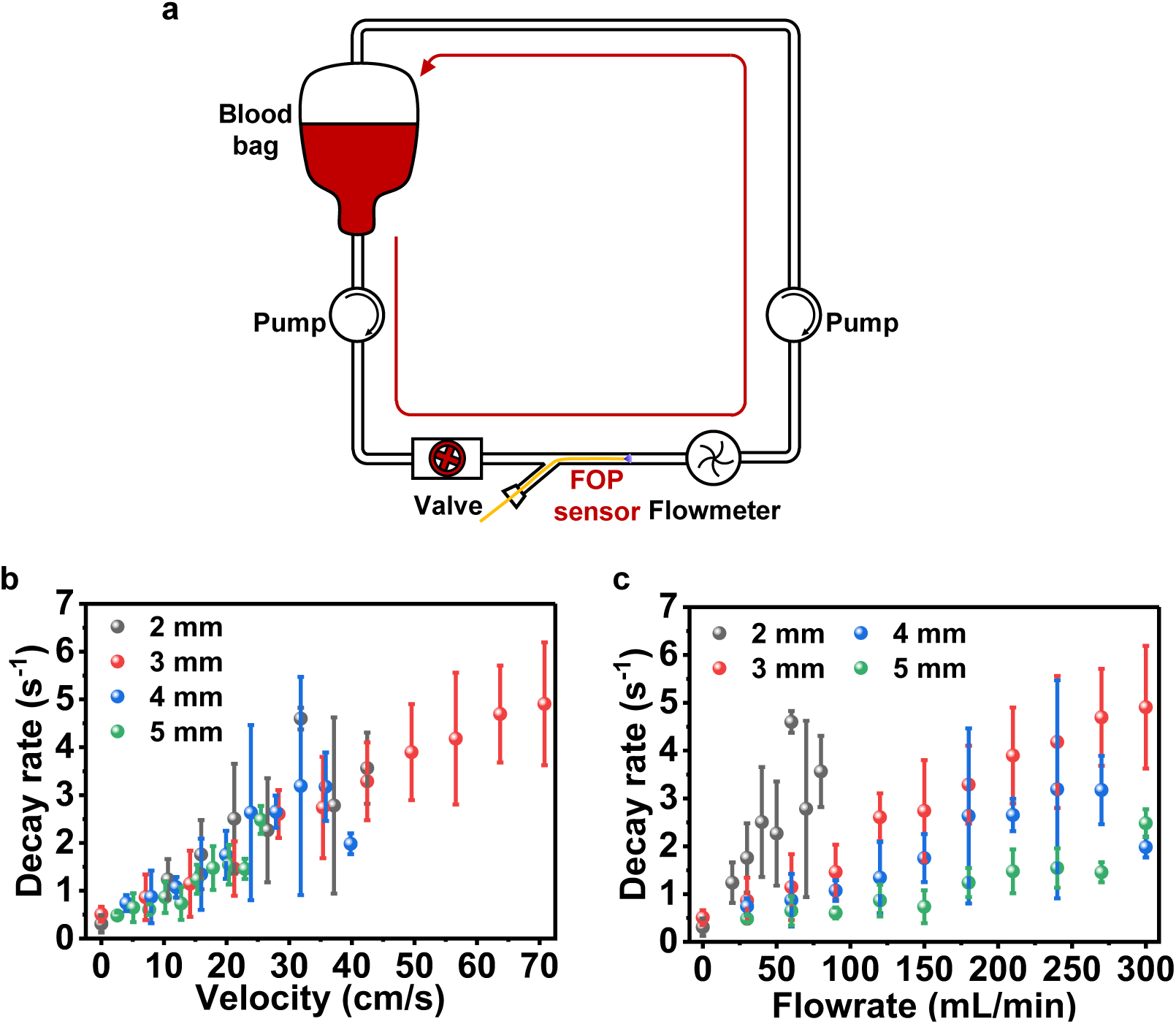
Blood velocimetry calibration. **a.** A customized in vitro blood circulation system used for calibration. **b, c.** Measured decay rate results obtained in tubes of different diameters: decay rate-velocity relationship **(b)** and decay rate-flowrate relationship **(c)**. These results indicate that the decay rate is linearly proportional to the flow velocity independent of tube diameters.

**Figure S18.**
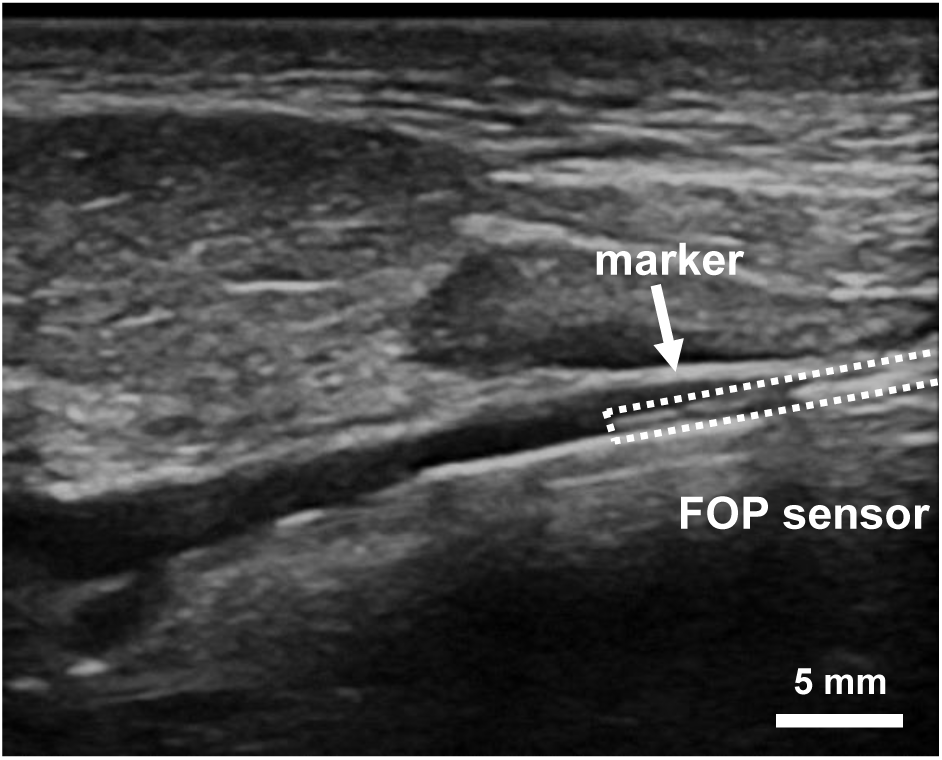
Ultrasonic image of the FOP sensor in the canine internal carotid artery.

**Figure S19.**
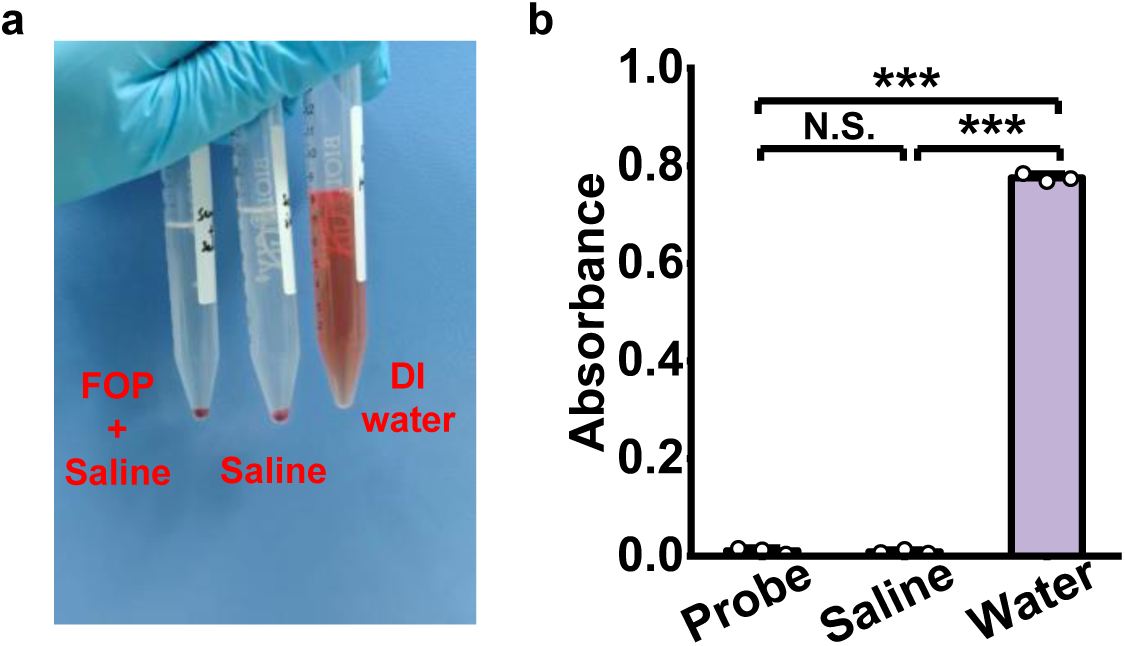
Hemolysis rate tests. **(a)** diluted blood incubated in different environments. **(b)** Absorbance at the wavelength of 545 nm of supernatant of these samples.

**Table S1.**
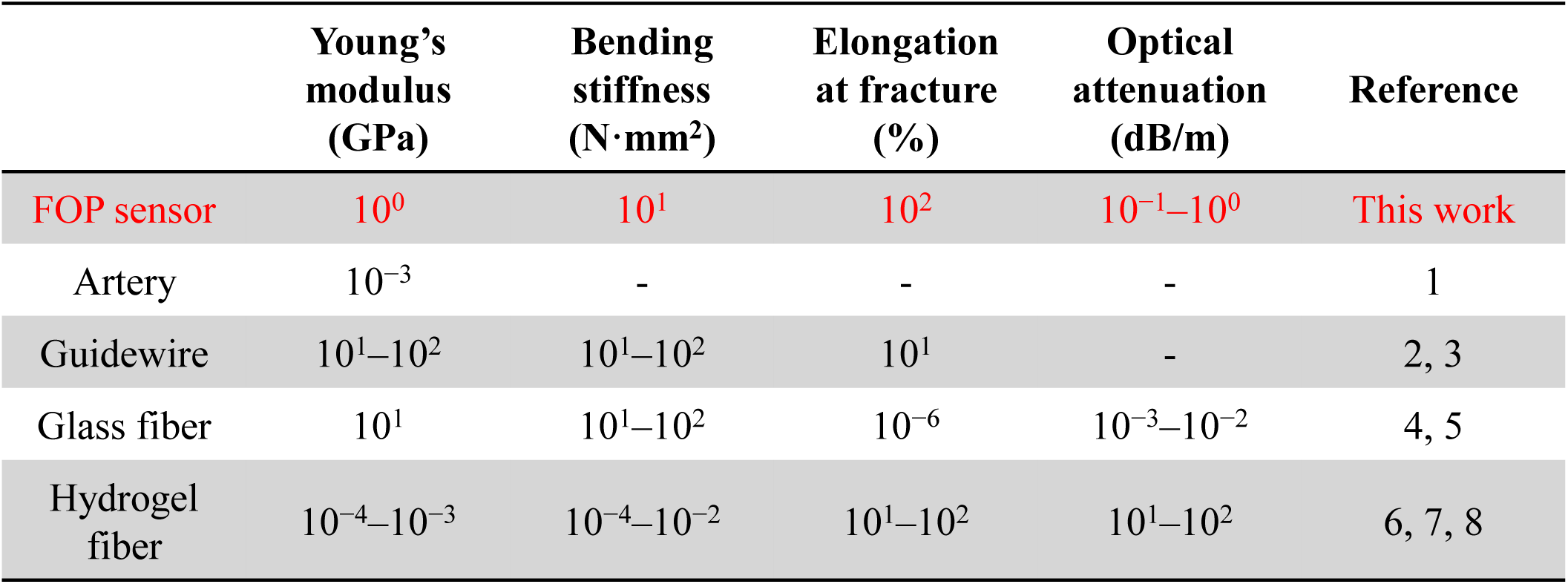
Comparison of mechanical and optical parameters among different sensors and tissues. Comparison between optical fiber used in FOP sensor, artery vessel, nitinol-based guidewire, glass and hydrogel based optical fibers.

|  | **Young’s modulus (GPa)** | **Bending stiffness (N·mm<sup>2</sup>)** | **Elongation at fracture (%)** | **Optical attenuation (dB/m)** | **Reference** |
|---|---|---|---|---|---|
| FOP sensor | 10<sup>0</sup> | 10<sup>1</sup> | 10<sup>2</sup> | 10<sup>−1</sup>–10<sup>0</sup> | This work |
| Artery | 10<sup>−3</sup> | - | - | - | 1 |
| Guidewire | 10<sup>1</sup>–10<sup>2</sup> | 10<sup>1</sup>–10<sup>2</sup> | 10<sup>1</sup> | - | 2, 3 |
| Glass fiber | 10<sup>1</sup> | 10<sup>1</sup>–10<sup>2</sup> | 10<sup>−6</sup> | 10<sup>−3</sup>–10<sup>−2</sup> | 4, 5 |
| Hydrogel fiber | 10<sup>−4</sup>–10<sup>−3</sup> | 10<sup>−4</sup>–10<sup>−2</sup> | 10<sup>1</sup>–10<sup>2</sup> | 10<sup>1</sup>–10<sup>2</sup> | 6, 7, 8 |

**Table S2.**
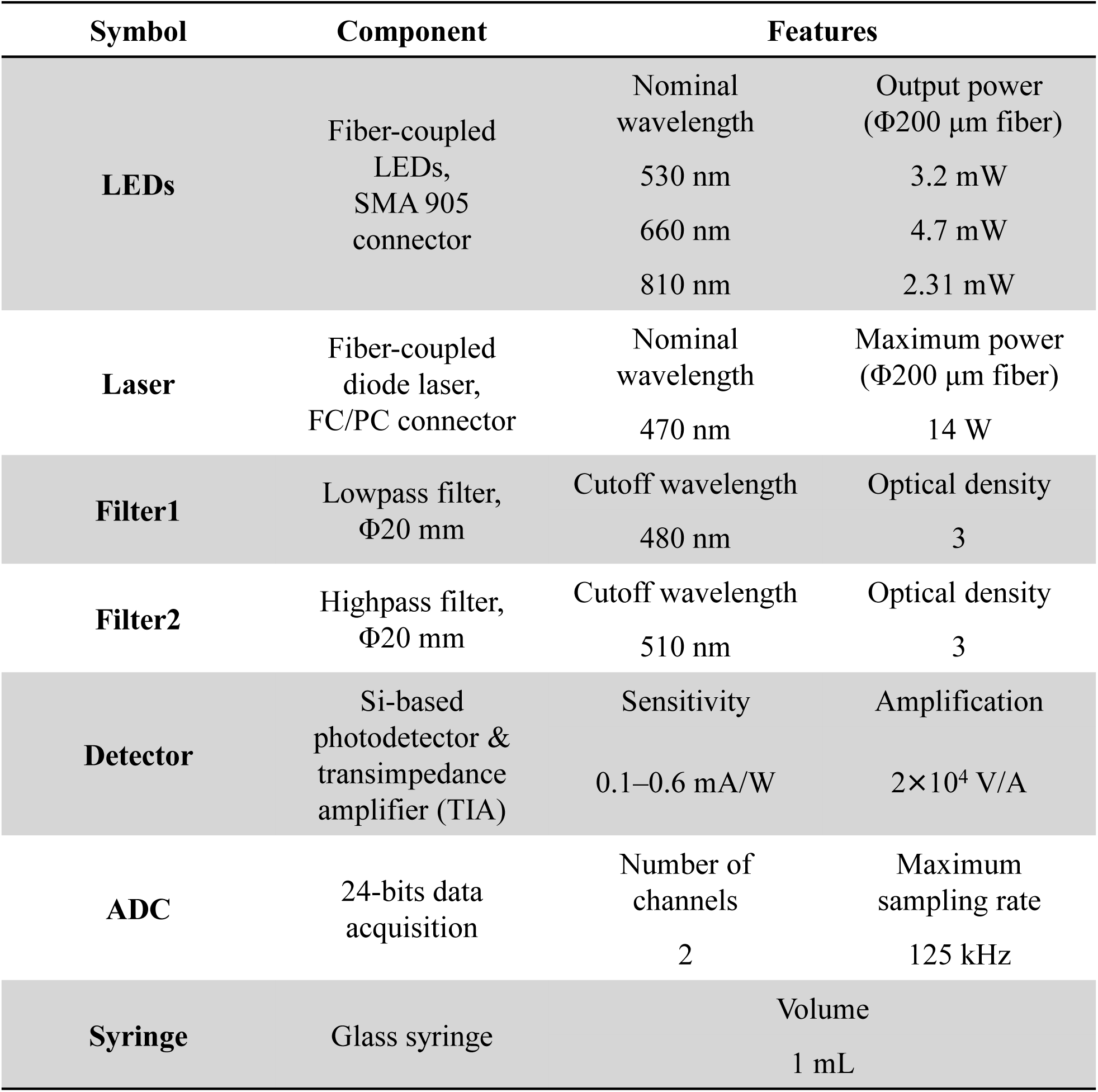
List of optical and fluidic components used in the FOP sensing system.

**Table S3.** List of parameters used in optical simulations.

|  |  | **Blood<br>(*S*<sub>O2</sub> = 98%)** | **Blood<br>(*S*<sub>O2</sub> = 0%)** | **Blood vessel** | **Tissue<br>(gray matter)** |
|---|---|---|---|---|---|
| **Green<br>(530 nm)** | ***μ*<sub>a</sub><br>(mm<sup>−1</sup>)** | 15.98 | 15.29 | 0.93 | 0.0431 |
|  | ***μ*<sub>s</sub><br>(mm<sup>−1</sup>)** | 106.16 | 95.91 | 31.78 | 10.7217 |
|  | **anisotropy<br>factor (*g*)** | 0.99 | 0.99 | 0.84 | 0.8863 |
| **Red<br>(660 nm)** | ***μ*<sub>a</sub><br>(mm<sup>−1</sup>)** | 0.18 | 1.83 | 0.34 | 0.0205 |
|  | ***μ*<sub>s</sub><br>(mm<sup>−1</sup>)** | 94.13 | 83.09 | 24.22 | 8.8758 |
|  | **anisotropy<br>factor (*g*)** | 0.99 | 0.99 | 0.85 | 0.8935 |
| **IR<br>(810 nm)** | ***μ*<sub>a</sub><br>(mm<sup>−1</sup>)** | 0.38 | 0.51 | 0.24 | 0.027 |
|  | ***μ*<sub>s</sub><br>(mm<sup>−1</sup>)** | 70.61 | 62.02 | 19.28 | 7.4853 |
|  | **anisotropy<br>factor (*g*)** | 0.98 | 0.98 | 0.86 | 0.8967 |

**Table S4.** List of parameters used in fluidic simulations.

|  | **Blood** | **Dye** |
|---|---|---|
| **Density<br>(kg/m<sup>3</sup>)** | 1060 | 993.36 |
| **Viscosity<br>(g/m·s)** | 0.00278 | 0.000692 |
| **Diffusion coefficient<br>(m<sup>2</sup>/s)** | 1.46×10<sup>−9</sup> |  |

**Table S5.** Comparing performance parameters (size, functionality and in-vivo monitoring location) of our FOP sensor and previously reported endovascular sensors.

| **Reference** | **Sensing modality** | **Operational principle** | **Subjects & Monitoring location** | **Probe diameter (mm)** |
|---|---|---|---|---|
| This work | Oxygen saturation | Optics | Rabbit femoral artery & Canine carotid artery | 0.5 |
|  | Flow velocity | Fluorescence dilution |  |  |
| 1 | Electro-physiology | Electrodes | Human superior sagittal sinus | 0.4 |
| 2 | Imaging | Optics | Human middle cerebral artery | 0.7 |
| 3 | Oxygen saturation | Optics | Human superior vena cava | 0.67 |
| 4 | Imaging | Ultrasonics | Human coronary artery | 1.15 |
|  | Flow velocity |  |  |  |
| 5 | Biochemical analytes | Electrochemistry | Rabbit<br>renal vein | 2 |
| 6 | Pressure | Capacitive | Porcine femoral artery | 0.8 |
| 7 | Temperature | Resistive | Porcine pulmonary artery | 3.7 |
|  | Pressure | Resistive |  |  |
|  | Flow velocity | Resistive |  |  |
| 8 | Pressure | Capacitive | Bypass graft at porcine carotid artery | 4 |

## Supporting Videos

**Movie S1.** An FOP sensor injecting a droplet of sodium fluorescein dye solution.

**Movie S2.** An FOP sensor passing through a bifurcating tube. The sensor microchannel is filled with ink for visualization.

**Movie S3.** An FOP sensor inserted through a rabbit artery.

**Movie S4.** Video of dye injection and dilution process (false-color) in slow (left, 13 cm/s) and fast (right, 53 cm/s) aqueous flow velocity.

**Movie S5.** Numerical simulation of dye dilution process in slow (left, 14 cm/s) and fast (right, 56 cm/s) blood flow.

**Movie S6.** An FOP sensor passing through vasculature into carotid arteries in the canine model. Video is captured by the DSA navigation system.

## Supporting Analysis

**Analysis S1.** Theoretical analysis on the signal processing principle of blood oxygenation calculation.

**Analysis S2.** Theoretical analysis on the signal processing principle of blood flow velocity calculation.

## Analysis S1. Method for Calculating Blood Oxygen Saturation

### 1. Relationship between recorded intensity and concentration of hemoglobins

The main target of data processing in oximetry is extracting blood oxygenation information from light intensity signals. The optical procedure of FOP oximetry includes two main parts: the extinction loss within the blood vessel and the transmission loss within the external systems.

The output light intensity *I* can be expressed as:

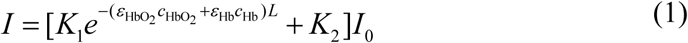

Here *I*_0_ is the incident light intensity, which is represented by the recorded signal at rest state at each wavelength (530, 660 and 810 nm). The first component is related to the extinction loss within the blood vessel, based on the Beer-Lambert Law^1^. *L* is optical path length for light travels in the blood at different wavelengths, which is determined by optical simulation. *ε*_HbO2_ and *ε*_Hb_ are molar extinction coefficients of oxygenated and deoxygenated hemoglobin (HbO_2_ and Hb), and *c*_HbO2_ and *c*_Hb_ are their corresponding concentrations. *ε*_HbO2_ and *ε*_Hb_ are determined according to literature and power spectral distribution of our light sources*. K*_1_ is a parameter related to the extinction of other components in blood, like water and other proteins^2^. *K*_2_ is related to the transmission loss within the external optic systems (fiber coupling, free space propagation, etc.). *K*_1_ and *K*_2_ are determined from in vivo BGA-based calibration in rabbit model.

We define the optical density (OD),

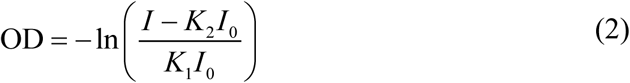

and get,

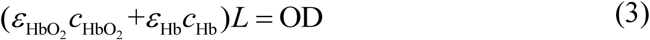

Combining equations of different wavelengths, we have an overdetermined system of linear equations,

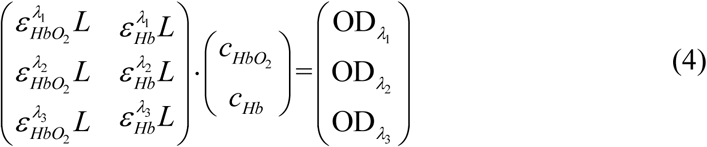

*c*_HbO2_ and *c*_Hb_ are solved from this system by least square method, from which blood oxygen saturation SO_2_ is determined,

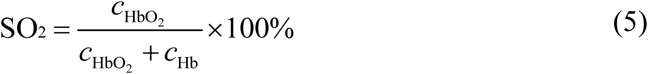

### 2. Determination of parameters

*ε*_HbO2_ and *ε*_Hb_ at different wavelengths used in data processing are extracted from literature^3–4^, and corrected based on the power spectral distribution of our LEDs. Because blood is a high-scattering media, *ε*_HbO2_ and *ε*_Hb_ used here are effective molar extinction coefficients, in which scattering is considered^3^. The coefficient is determined by:

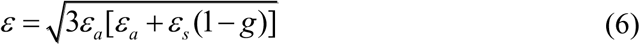

*ε*_a_ is molar absorption coefficient, *ε*_s_ is molar scattering coefficient, and *g* is scattering anisotropy. The value of *ε* is further corrected based on the power spectral density distribution of our light sources:

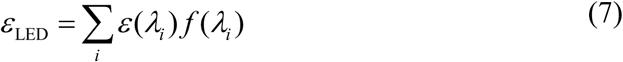

*f* is the power distribution function of LED light source across the spectrum, which is measured by a spectroscope. *ε*_LED_ is eventually used in SO_2_ calculation.

*L* at different wavelengths is determined from Monte Carlo simulation (Tracepro free trial version). In the model, a disc light source is placed in a block of blood with sufficient size, and the path length of every trace travelling back to the source is recorded. The average travelling distance of these traces is used as *L* in data processing.

*K*_1_ and *K*_2_ are determined from in vivo BGA-based calibration, where groups of pairing data of *c*_HbO2_, *c*_Hb_, and *I* are obtained. OD is calculated based on *ε*_HbO2_, *ε*_HbO2_, *L*, *c*_HbO2_, and *c*_Hb_, from which *K*_1_ and *K*_2_ were determined by parameter optimization.

The parameters used in parameter determination and blood oxygen saturation calculation are listed in Table 1.

**Table.1.** Parameters used in oximetry data processing.

|  | Wavelength |  |  |
|---|---|---|---|
|  | 530 nm | 660 nm | 810 nm |
| *ε*<sub>LED-HbO2</sub> (g<sup>−1</sup>·dL·mm<sup>−1</sup>) | 2.00 | 0.0691 | 0.100 |
| *ε*<sub>LED-Hb</sub> (g<sup>−1</sup>·dL·mm<sup>−1</sup>) | 2.19 | 0.310 | 0.102 |
| *K*<sub>1</sub> | 0.222 | 0.121 | 0.323 |
| *K*<sub>2</sub> | 0.969 | 0.939 | 0.841 |
| *L* (mm) | 0.0746 | 0.703 | 0.534 |

## Analysis S2. Method for Calculating Blood velocity

The main target of this analysis is to model the decay curve of fluorescent intensity and construct an algorithm for calculating blood velocity based on curves recorded by our FOP sensor implanted in the blood vessel. The intensity curve is determined by dispersion of dye molecules in the blood flow, as well as optical interaction between light, dye molecules, and blood. These processes will be discussed in this analysis.

### 1. Distribution of dye

Since the blood vessel has a high aspect ratio, and blood flows along the vessel, longitudinal procedures dominate in dye dispersion. As the blood vessel is a channel with a high aspect ratio, the axial processes dominate in the mass transfer of dye, one-dimensional model is implemented in this analysis^1^.

At the beginning of every velocimetry process, a bolus of dye solution is injected into blood flow. We assume that the concentration of dye in blood follows Gaussian distribution upon injection^2^,

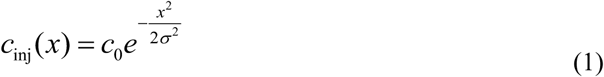

*c*_inj_(*x*) is the concentration of dye at location *x*, *c*_0_ is related to the total amount of injected dye, *σ* describes the distribution deviation of a bolus. Under strict control of injection pump, *c*_0_ and *σ* are considered as constants for every trial.

This bolus of dye subsequently experiences dispersion in blood flow, which could be described by the convection equation,

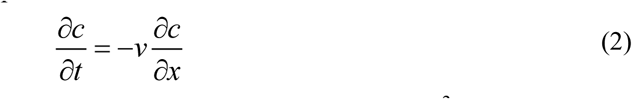

The Péclet Number (*Pe*) of this process is defined as *Pe* = *vL*/*D*^3^, where *v* is the velocity of fluid, *L* is the length of the region of interest, *D* is the diffusion coefficient of dye molecule. Here *L* is ∼ mm scale in endovascular circumstance, *D* is ∼ 10^−4^ mm^2^/s for an aromatic organic small-molecule dye anion^4^. In the normal range of arterial blood velocity (20–200 cm/s)^[5]^, *Pe* ∼ 10^6^–10^7^, which is much larger than 1, indicating that dye diffusion can be neglected in this process^3^. Therefore, only fluidic transport and convention is considered here.

To solve equation (2), spatial distribution (1) is used as initial condition. The solution shows the spatial-temporal distribution of dye concentration in blood flow,

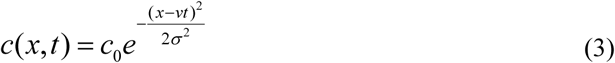

### 2. Fluorescent photometry

The fluorescent intensity *I* recorded by the FOP sensor is determined by incident light intensity from fiber, optical extinction of blood, as well as concentration distribution, absorption coefficient, and quantum yield of the dye^6^. Thus, the fluorescent intensity *I* could be depicted as,

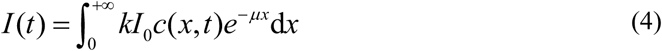

*k* is a parameter related to the quantum yield and the absorption coefficient at the stimulating wavelength of the fluorescent dye, *I*_0_ is the incident light entering blood, *μ* is related to extinction coefficient of blood at stimulating and fluorescent wavelengths. Due to stable optical properties of dye and blood during the trials, *k* and *μ* are regarded as constants.

Combining expression (3) and (4), and we get,

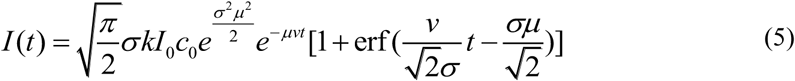

This expression depicts the recorded fluorescent intensity evolution after a typical injection.

### 3. Blood velocity calculation based on curve feature

Due to the complex form of expression (5), it is not practical to directly fit the recorded intensity curve with this equation to obtain *v*. Therefore, we chose a curve feature with a linear relationship with *v* as the indicator of blood velocity. The chosen indicator is the normalized maximum decreasing rate of the intensity curve, the ratio between the absolute value of the minimum of derivative of the curve and the maximum of the curve, abs[min(d*I*/d*t*)]/max(*I*). This term is proportional to the blood velocity *v*, which is proved as follow.

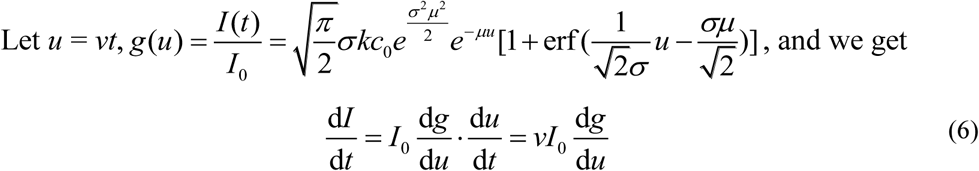

Since *v* and *I*_0_ are independent of *t* during each trial, the minimum point of d*I*/d*t* (*t*|_min(d*I*/d*t*)_) corresponds to the minimum point of d*g*/d*u* (*u*|_min(d*g*/d*u*)_). So, we get

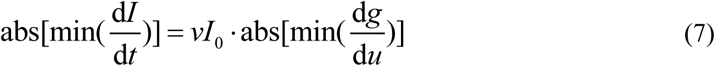

Because *v* and *I*_0_ is not present in function *g*(*u*), abs[min(d*g*/d*u*)] is a constant across trials, determined by constants *σ*, *k*, *c*_0_, and *μ* only. Therefore, abs[min(dI/dt)] is proportional to *v* and *I*_0_.

The next step is to remove the influence of *I*_0_ from the indicator, for the incident light intensity from the laser source drifts across trials.

The maximum intensity max(*I*) of the curve is chosen to represent *I*_0_. The reason is that *v* is not present in *I*_0_·*g*(*u*) (always equal to *I*(*t*)), so that its maximum is proportional to *I*_0_ and non-relevant to *v*:

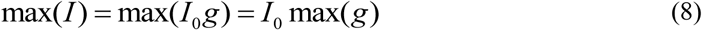

Here max(*g*) is determined by constants *σ*, *k*, *c*_0_, and *μ* only. By combining expression (7) and (8), we remove the influence of *I*_0_ and get the linear indicator of velocity used in FOP velocimetry, abs[min(d*I*/d*t*)]/max(*I*), which is defined as the decay rate,

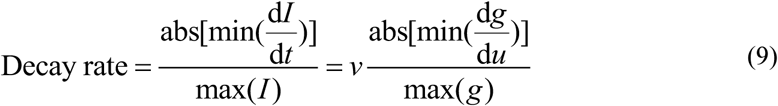

The proportional coefficient between decay rate and *v* (abs[min(d*g*/d*u*)]/max(*g*)) is determined by optical and fluid dynamics properties of blood and dye solution, and independent of *v* and *I*_0_. In a typical FOP velocimetry trial, abs[min(d*I*/d*t*)] and max(*I*) are extracted from the intensity decay curve, and the indicator decay rate is calculated. Blood velocity is further obtained from this indicator.

The proportional coefficient between *v* and decay rate used in velocity calculation is obtained through calibration experiments (Fig. 3e).

